# Measuring glycolytic flux in single yeast cells with an orthogonal synthetic biosensor

**DOI:** 10.1101/682302

**Authors:** Francisca Monteiro, Georg Hubmann, Justin Norder, Johan Hekelaar, Joana Saldida, Athanasios Litsios, Hein J. Wijma, Alexander Schmidt, Matthias Heinemann

**Author notes:** cE3c-Centre for Ecology, Evolution and Environmental Changes, Faculdade de Ciências, Universidade de Lisboa, Campo Grande, 1749-016, Lisboa, Portugal (Francisca Monteiro). Department of Biology, Laboratory of Molecular Cell Biology, Institute of Botany and Microbiology, KU Leuven, & Center for Microbiology, VIB, Kasteelpark Arenberg, 31, 3001 Heverlee, Flanders, Belgium. Shared first authors (Francisca Monteiro, Georg Hubmann).

## Abstract

Metabolic heterogeneity between individual cells of a population harbors offers significant challenges for fundamental and applied research. Identifying metabolic heterogeneity and investigating its emergence requires tools to zoom into metabolism of individual cells. While methods exist to measure metabolite levels in single cells, we lack capability to measure metabolic flux, i.e. the ultimate functional output of metabolic activity, on the single-cell level. Here, combining promoter engineering, computational protein design, biochemical methods, proteomics and metabolomics, we developed a biosensor to measure glycolytic flux in single yeast cells, by drawing on the robust cell-intrinsic correlation between glycolytic flux and levels of fructose-1,6-bisphosphate (FBP), and by transplanting the *B. subtilis* FBP-binding transcription factor CggR into yeast. As proof of principle, using fluorescence microscopy, we applied the sensor to identify metabolic subpopulations in yeast cultures. We anticipate that our biosensor will become a valuable tool to identify and study metabolic heterogeneity in cell populations.

## Introduction

Increasing evidence suggests that individual cells in a population can be metabolically very different^1–5^. Metabolic heterogeneity has been found, for instance, in microbial cultures used for biotechnological processes^6^, but also in cells of human tumours^7^. Because metabolic heterogeneity is connected with productivity and yield losses in biotechnological production processes^6^, and in cancer with limited therapeutic successes^8^, it is key to identify metabolic subpopulations and to understand their emergence.

Towards assessing metabolic heterogeneity, several novel experimental tools have recently been developed to measure metabolite levels in single cells, e.g. by exploiting the autofluorescence of specific metabolites^9^, Förster resonance energy transfer (FRET)^10^, or metabolite-binding transcription factors^11^. For instance, transcription factor (TF)-based biosensors now exist to detect amino acids^12^, sugars^13^ and the metabolites succinate and 1-butanol^14^, tricacetic acid lactone^15^ or malonyl CoA^16^, partly enabled by the transplantation of prokaryotic metabolite-responsive TFs to eukaryotes^17–21^.

While measurements of metabolite *levels* in single cells are already useful, knowledge of metabolic *fluxes* in individual cells would often be more informative, as metabolic fluxes represent the ultimate functional output of metabolism. Therefore, they serve as predictor of productivity in the development of cell factories^22^, or as indicator of disease^23^. Here, particularly knowing the flux through glycolysis would be valuable, as this flux has been shown to correlate with highly-productive phenotypes^24^ and cancer^25^. While nowadays metabolic fluxes can be resolved in ensembles of cells, for instance, by means of ^13^C-metabolic flux analysis^26^, inference of fluxes in individual cells, however, is so far not possible, but yet highly desired^5^.

One possible avenue towards measuring metabolic fluxes in individual cells has recently emerged by the discovery of so-called flux-signalling metabolites^27^, which are metabolites, whose levels - by means of particular regulation mechanisms^28^ - strictly correlate with the flux through the respective metabolic pathway. Such flux signals are used by cells to perform flux-dependent regulation^29, 30^. Biosensors for such metabolites, such as recently accomplished for *E. coli*^31^, would in principle allow for measurement of metabolic fluxes in single cells, in combination with microscopy or flow cytometry.

Here, drawing on glycolytic flux-signalling metabolite fructose-1,6-bisphosphate (FBP) levels in yeast^30^ and using the *B. subtilis* FBP-binding transcription factor CggR^32, 33^, we developed a biosensor, which allows to measure glycolytic flux in single yeast cells. To this end, we used computational protein design, biochemical, proteome and metabolome analyses, for (i) the development of a synthetic yeast promoter regulated by the bacterial transcriptional factor CggR, (ii) the engineering of the transcription factors’ FBP binding site towards increasing the sensor’s dynamic range, and (iii) the establishment of growth-independent CggR expression levels. We demonstrate the applicability of the biosensor for flow cytometry and time-lapse fluorescence microscopy. We envision that the biosensor will open new avenues for both fundamental and applied metabolic research, not only for monitoring glycolytic flux, but also for engineering control circuits with glycolytic flux as input variable.

## Results

### Design of biosensor concept

For our biosensor, we exploited the fact that the level of the glycolytic intermediate fructose-1,6-biphosphate (FBP) in yeast^30, 34^, similar to other organisms^27^, strongly correlates with the glycolysis flux^28, 34, 35^, and that changing FBP levels exert flux-dependent regulation. In *B. subtilis*, for instance, FBP binds to the transcription factor (TF) CggR^33^, which when bound to its target DNA forms a tetrameric assembly of two dimers, through which transcription gets inhibited^36^. Upon binding of FBP to the CggR-DNA complex, the dimer-dimer contacts of CggR are disrupted, and the promoter is derepressed^37^.

Here, we aimed to transplant the *B. subtilis* CggR to yeast and have it exerting FBP-, and thus, glycolytic flux-dependent expression of a fluorescent protein. To this end, a number of challenges had to be addressed. First, a synthetic promoter had to be designed for the foreign transcription factor CggR, involving the identification of ideal positioning and number of operator sequences^38, 39^, and engineering the nucleosome architecture to allow for maximal promoter activity^40^. Second, CggR had to be made responsive to FBP in the correct dynamic range, requiring protein engineering efforts^13, 41^. Third, the CggR expression levels needed to be such that together with the metabolite-modulating effect of CggR, the TF can actually exert a regulating effect on the promoter in the right range, for which we needed to identify the right CggR expression levels (Figure 1).

**Figure 1.**
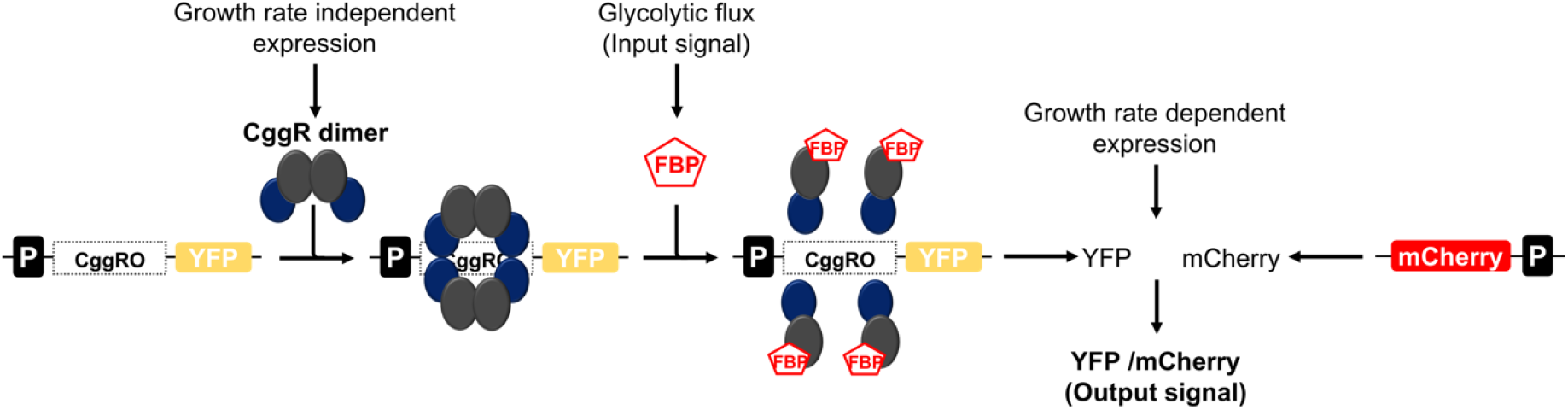
Illustration of the biosensor concept to measure glycolytic fluxes in single *S. cerevisiae* cells. Expression of the bacterial transcriptional repressor CggR at constant levels, i.e. independent of growth rate and substrates. Binding of CggR as a dimer of dimers to the operator (CggRO) of the synthetic cis-regulatory region, forming the CggR-DNA complex repressing transcription. At high glycolytic fluxes, fructose-1,6-bisphosphate (FBP) levels are high and FBP binds to CggR disrupting the dimer-dimer contacts, which induces a conformational change of the repressor, such that transcription of the reporter gene (YFP) can occur. The binding of FBP to CggR and consequent transcription is dependent on the FBP concentration, which is correlated with the glycolytic flux. The activity of the glycolytic flux biosensor is measured by quantifying YFP expression. YFP expression levels are normalized through a second reporter, mCherry, under the control of TEF1 mutant 8 promoter (P_TEFmut8_), to control for global variation in protein expression activity.

### *In vivo* test system for a substrate- and growth rate-independent flux-sensor

For later evaluation of the flux-reporting capacity of the developed sensor, we first established an *in vivo* test system, through which we could generate a range of glycolytic fluxes at steady-state conditions. To this end, we employed a combination of growth substrates and two different *S. cerevisiae* strains: the wildtype (WT) and a mutant strain (TM6), which only carries a single chimeric hexose transporter and thereby only generates low glucose uptake rates at high glucose levels^42^. Metabolome and physiological analyses in combination with a new method for intracellular flux determination^43^ showed that this combination of strains and conditions allowed us to generate a broad range of glycolytic fluxes (Figure 2A). Consistent with the earlier reported correlation between FBP levels and glycolytic flux^30^, also here the FBP levels linearly correlated with flux (R^2^ = 0.99; p<0.0001) (Figure 2A), but not with growth rate (Figure 2B). This set of conditions and strains, thus served as test system for the to-be developed glycolytic flux sensor.

**Figure 2.**
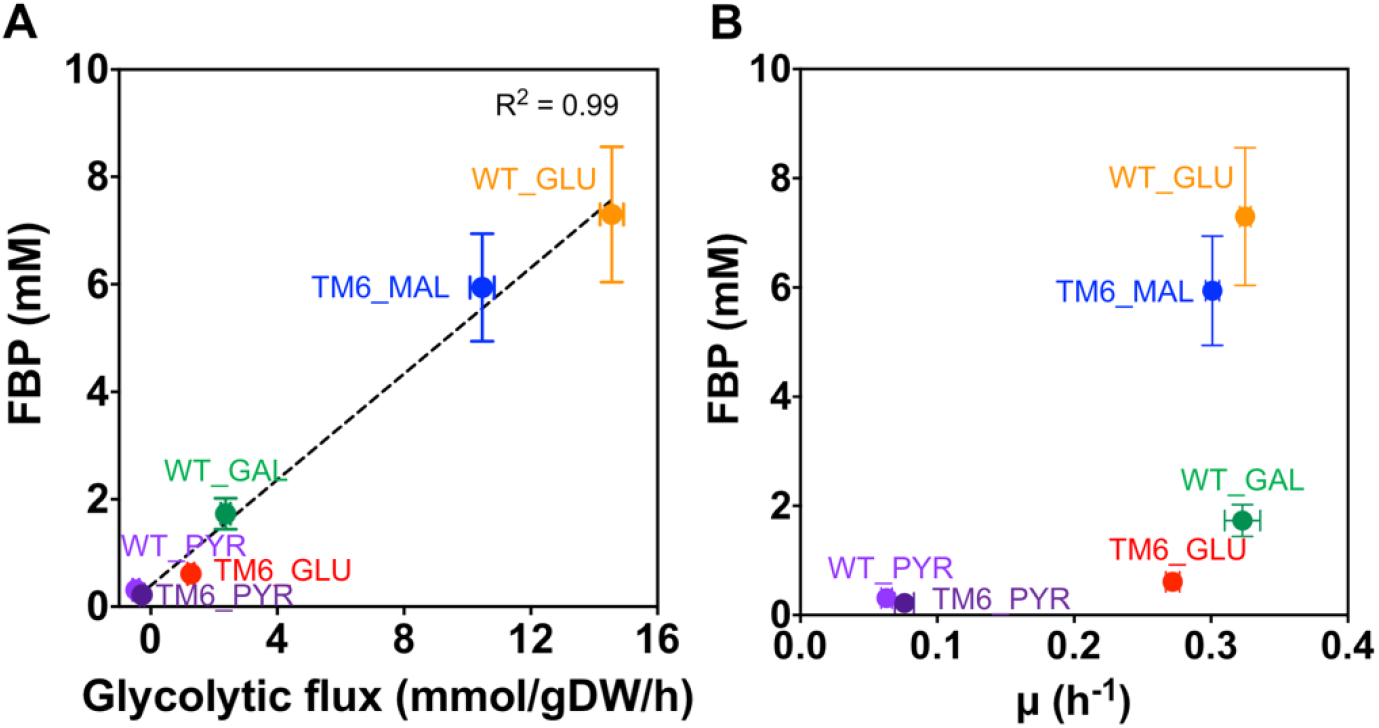
FBP concentration linearly correlates with glycolytic flux, but not with growth rate. (A) Glycolytic flux of wildtype (WT) and TM6 strains correlates with fructose-1,6-bisphosphate (FBP) concentration. The glycolytic flux is here reported as the flux between the metabolites fructose 6-phosphate (F6P) and FBP. Glycolytic fluxes were here estimated on the basis of physiological and metabolome data and a novel method to estimate intracellular fluxes^43^. While on high glucose, the WT strain accomplishes a high glucose uptake rate (and thus glycolytic flux), the mutant strain (TM6) only generates a low glucose uptake (and thus glycolytic flux). On maltose, also the mutant strain achieves a high glycolytic flux, since maltose is transported via a separate transporter^44^. (B) FBP concentration as a function of cellular growth rate shows no correlation. For metabolite levels and growth rates, error bars correspond to the standard deviation between three independent experiments; for glycolytic fluxes to the mean and standard deviations of the sampled flux solution space (cf. Methods). The carbon sources were used at a final concentration of 10 gL^-1^ and are indicated: Glucose (GLU); Galactose (GAL); Maltose (MAL) and Pyruvate (PYR)

### Development of the synthetic CggR cis-regulatory element

First, we designed a synthetic CggR cis-regulatory element for yeast (CggRO) based on the *CYC1* promoter, which was previously successfully re-designed^40^. To accomplish repression of the promoter by CggR, we aimed to shield the TATA boxes by the binding and tetramerization of the CggR dimers. The *CYC1* core promoter has three TATA boxes at the positions −221, −169, and −117, upstream of the open reading frame (Figure 3A –upper part). We flanked the two TATA boxes at position −221 and −117 up- and downstream with a CggR operator site. To conserve the geometry of the *CYC1* core promoter as much as possible, we removed the TATA box at position −169, because this TATA box was exactly located, where we integrated the CggR operator sites flanking the other TATA boxes, and we did not want to make the sequence longer. The 5’UTR of the *CYC1* promoter, which also included the transcriptional start site, was kept. To allow for sole binding and regulation through CggR, we removed the part further upstream of the TATA box at the position −221, were according to YEASTRACT^45^, the endogenous transcriptional binding sites of the *CYC1* promoter are located.

**Figure 3.**
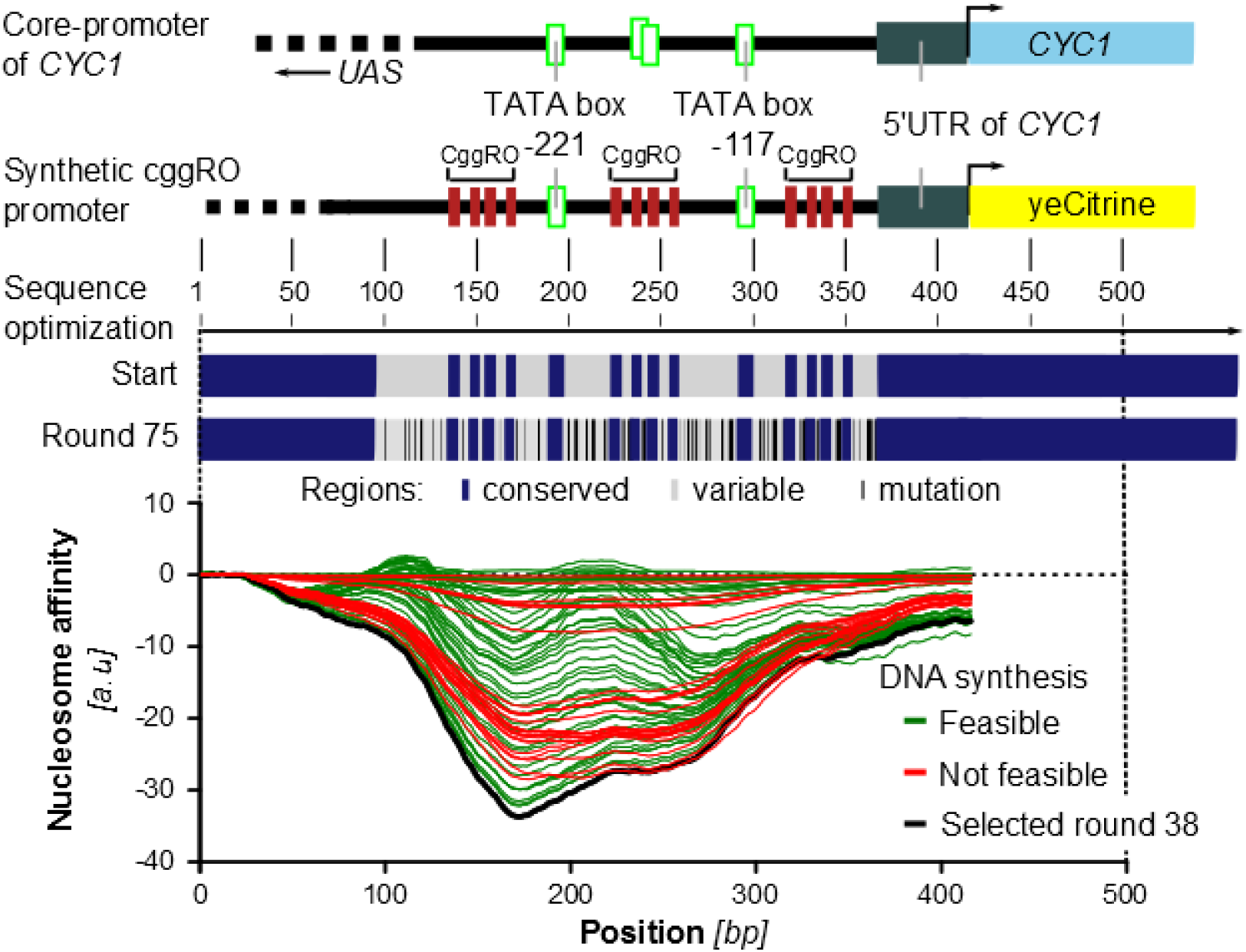
Design of the synthetic CggR cis-regulatory element. The promoter design is based on the *CYC1* core promoter. The relevant structural elements of the *CYC1* core promoter elements, which are required for transcription, were conserved in the synthetic promoter design. These elements comprised two TATA boxes at position –221 and –117 (relative to the start of the *CYC1* ORF), and the 5‘UTR of the *CYC1* core promoter (including transcriptional start site, TSS). In the promoter design, three CggR operator sites were inserted adjacent to the two TATA boxes. All functional elements were conserved (blue colored region) during the optimization of the promoter sequence. Nucleotide sequences between the functional elements (grey colored region) were allowed to be optimized by the algorithm. Nucleotides that got optimized are indicated with a black line. A total of 75 sequence versions were generated, where each sequence differed in one mutation from the progenitor sequence. The sequences were optimized for low nucleosome affinity. After optimization, all sequences were checked for synthesis feasibility. The synthesis of the sequences was feasible (green) until the 46^th^ round. After this round, the sequences (not feasible in red) reached a GC content insufficient for proper synthesis. The promoter sequence, which was generated in round 38 (black), showed the best compromise between nucleosome affinity and the possibility to synthesize the sequence.

Using a recently published computational method^40^, we further optimized this designed sequence of the CggR cis-regulatory element to minimize nucleosome binding. Functional elements (e.g. the CggR operator sites, the TATA boxes and the 5’UTR; cf. Supplementary Table 1) were excluded from the sequence optimization (Figure 3 – lower part). 75 computational optimization rounds were applied. As the CggR cis-regulatory element resembled a repetitive DNA sequence with a high AT content, sequence variants were checked for DNA synthesis feasibility. The cis-regulatory element of round 38 was the variant with the lowest nucleosome affinity with retained feasibility for DNA synthesis. The synthesized synthetic promoter was integrated upstream of the fluorescent reporter protein YFP (eCitrine) in a centromeric plasmid ensuring a stable copy number.

### Establishing a substrate- and growth rate-independent CggR expression

Next, to drive expression of CggR, we needed a promoter that would lead to condition-independent (i.e. constant) intracellular CggR levels in order to ensure that the flux sensor only reports altered FBP levels (i.e. glycolytic fluxes), and not altered CggR levels. To this end, we tested the P_CMV_ promoter, which is widely used as a strong constitutive promoter in mammalian cells^46^, and two mutant variants of the endogenous TEF1 promoter, i.e. mutant 2 (P_TEFmut2_) with low, and mutant 7 (P_TEFmut7_) with medium-to-high expression strength^47^. Each promoter and the CggR gene were cloned into the HO genomic locus of both yeast strains.

To quantify the CggR protein levels, we performed proteome analyses with the different strains, promoters and growth conditions. Overall, the three promoters yielded largely different CggR abundances on glucose (Figure 4A). Across conditions and thus growth rates, we found that the CggR levels when expressed from the P_CMV_ and P_TEFmut2_ promoters showed significant variations, while the P_TEFmut7_ promoter generated more comparable levels across growth rates (Figure 4B), as established through the different carbon sources and strains. Because of its condition-independent expression level, we selected the P_TEFmut7_ promoter to drive the CggR expression.

**Figure 4.**
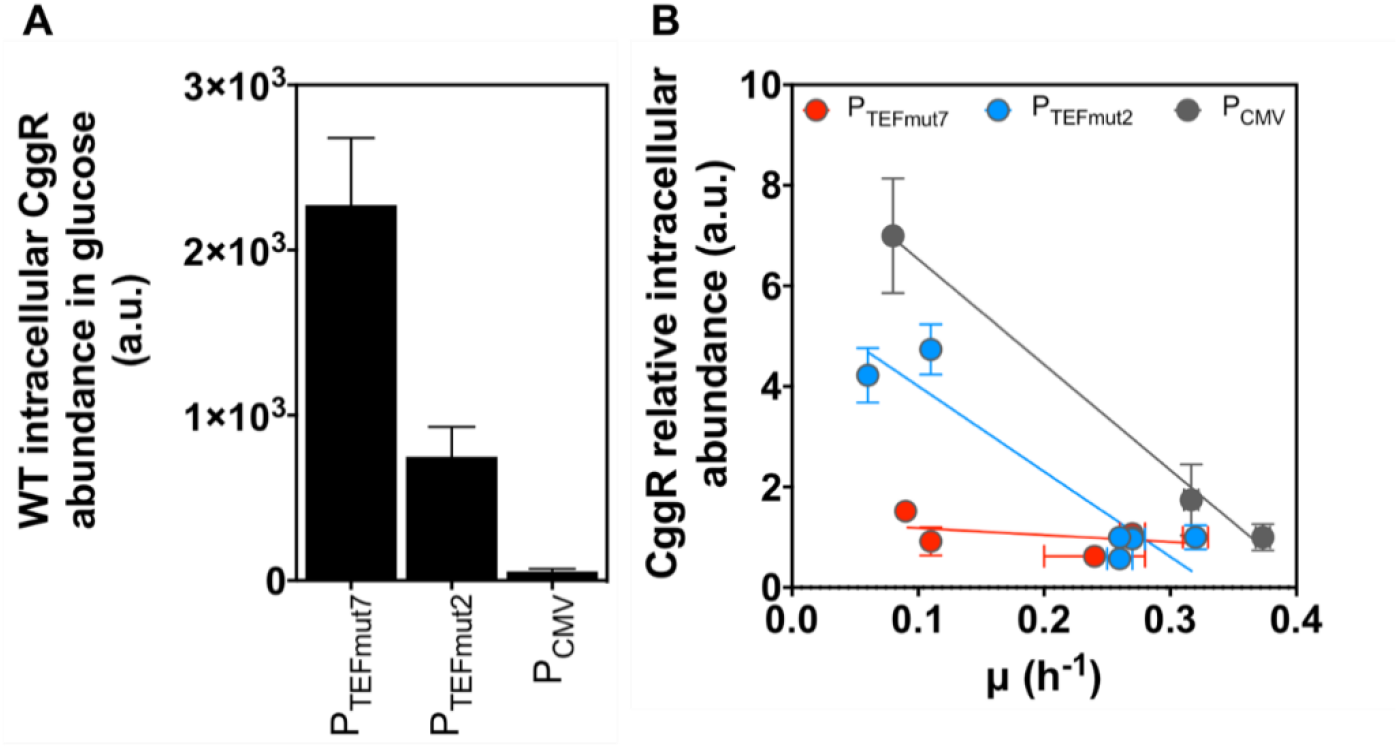
CggR intracellular levels and expression profile with different promoter, strains and conditions. (A) CggR intracellular abundance in the wildtype (WT) strain on glucose strongly varies with the promoter used. (B) The relative abundance of CggR (normalized to the abundance measured on glucose and the same promoter) is almost constant with P_TEFmut7_ across multiple growth rates, but not with P_TEFmut2_ and P_CMV_. The CggR intracellular levels were quantified by proteomics in steady-state cultures. WT data includes all three promoters whereas TM6 only includes the P_TEFmut2_ and P_TEFmut7_ data. Error bars represent the standard deviation of at least three replicate experiments.

### Engineering the FBP affinity of CggR

Next, we needed to engineer the FBP binding to CggR, such that this matches with the physiological range of FBP levels. FBP levels in yeast range from 0.2 mM to around 8 mM (Figure 2A). As the wildtype CggR has an affinity for FBP of around 1 mM^32^, we needed to generate a CggR mutant with a lower affinity for FBP, and with ideally a graded interaction between CggR and FBP towards accomplishing a broad linear dynamic range of the sensor. Importantly, the engineered CggR would still need to bind to the DNA, and furthermore the protein should be stable to not affect its cellular abundance.

To obtain such a CggR mutant, supported by computational protein design methods, we identified mutations at the FBP binding site that could lead to the desired decrease in affinity. Specifically, as in the CggR structure (3BXF)^48^ CggR binds to FBP through hydrogen bonds, we designed mutations to weaken or disrupt H-bonding interactions (Table 1, Supplementary Table 2), with the aim to decrease binding affinity. The X-ray structure further showed that FBP-binding causes a conformational change of CggR^48^, where a loop between residues G177 and Q185 moves away from the FBP-binding site towards another subunit. On the basis of this, we conjectured that mutations might not only influence FBP-binding, but might alter the equilibrium between the normal and activated conformation, even in absence of FBP. To predict the effect of the mutations on this equilibrium, and on overall protein stability, we used FoldX^49^, where we found that a E269Q mutation could decrease overall stability while R175K could permanently shift CggR to its activated conformation (Table 1). Four mutations (i.e. T151S, T151V, T152S, and R250A) were thus identified as promising candidates for decreasing the FBP-binding to CggR without otherwise negative effects (Table 1).

**Table 1.** List of predicted mutations to alter CggR-FBP binding affinity.

| Mutation | Expected effects on affinity for FBP (and if relevant on equilibrium and stability)<sup>a</sup> | FoldX predicted stability changes (kJ/mol)<sup>b</sup> |  |  |
| --- | --- | --- | --- | --- |
| | | $\Delta\Delta G^{\text{fold}}$ for the normal-conformation | $\Delta\Delta G^{\text{fold}}$ for the activated conformation | $\Delta\Delta\Delta G^{\text{fold}}$ (between the conformations) |
| **T151S** | **negligible to mild** affinity decrease | -1.4 | -1.3 | 0.1 |
| **T151V** | **mild to strong** affinity decrease | 2.5 | 3.1 | 0.5 |
| **T152S** | **negligible to mild** affinity decrease | 7.3 | 4.6 | -2.7 |
| **R175K** | **negligible to strong** affinity decrease and possibly a **shift of equilibrium** to the activated conformation | 16.5<sup>c</sup> | -0.7 | -17.2<sup>c</sup> |
| **R250A** | **mild to strong** affinity decrease | 2.2 | 2.7 | 0.5 |
| **E269Q** | **mild to strong** affinity decrease for FBP in combination with **overall destabilization** | 16.4<sup>c</sup> | 11.4<sup>c</sup> | -5.0 |

<sup>a</sup>A detailed justification for the expected effects of the mutations on binding affinity is given in Supplementary Table 2. <sup>b</sup>A downshift in $\Delta\Delta G^{\text{fold}}$ predicts stabilization of the protein, while a downshift in $\Delta\Delta\Delta G^{\text{fold}}$ predicts that the FBP-conformation becomes more favorable. $\Delta\Delta\Delta G^{\text{fold}}$ represents the difference between the $\Delta\Delta G^{\text{fold}}$ values for the two conformations. <sup>c</sup>Values are significantly higher than the standard deviation of FoldX predictions, which equals 3.4 kJ/mol<sup>49</sup>.

Next, we generated these CggR mutants with site-directed mutagenesis, expressed in *E. coli*, purified, and biochemically characterized them. To this end, we used thermal shift assays to assess protein stability and ligand binding. Most of the engineered CggR variants maintained wildtype stability, with the exception of E269Q (consistent with the above analysis) and T151V, which were less stable as indicated by decreased melting temperatures (Figure 5A). While the wildtype had a K_D_ of 1 mM FBP, the mutants T151S, E269Q and T152S showed a 1.1, 1.5 and 1.6-fold lower K_D_ values, respectively, while the K_D_ values of the R250A and T151V mutants increased 1.5 and 2.6-fold (Figure 5B).

**Figure 5.**
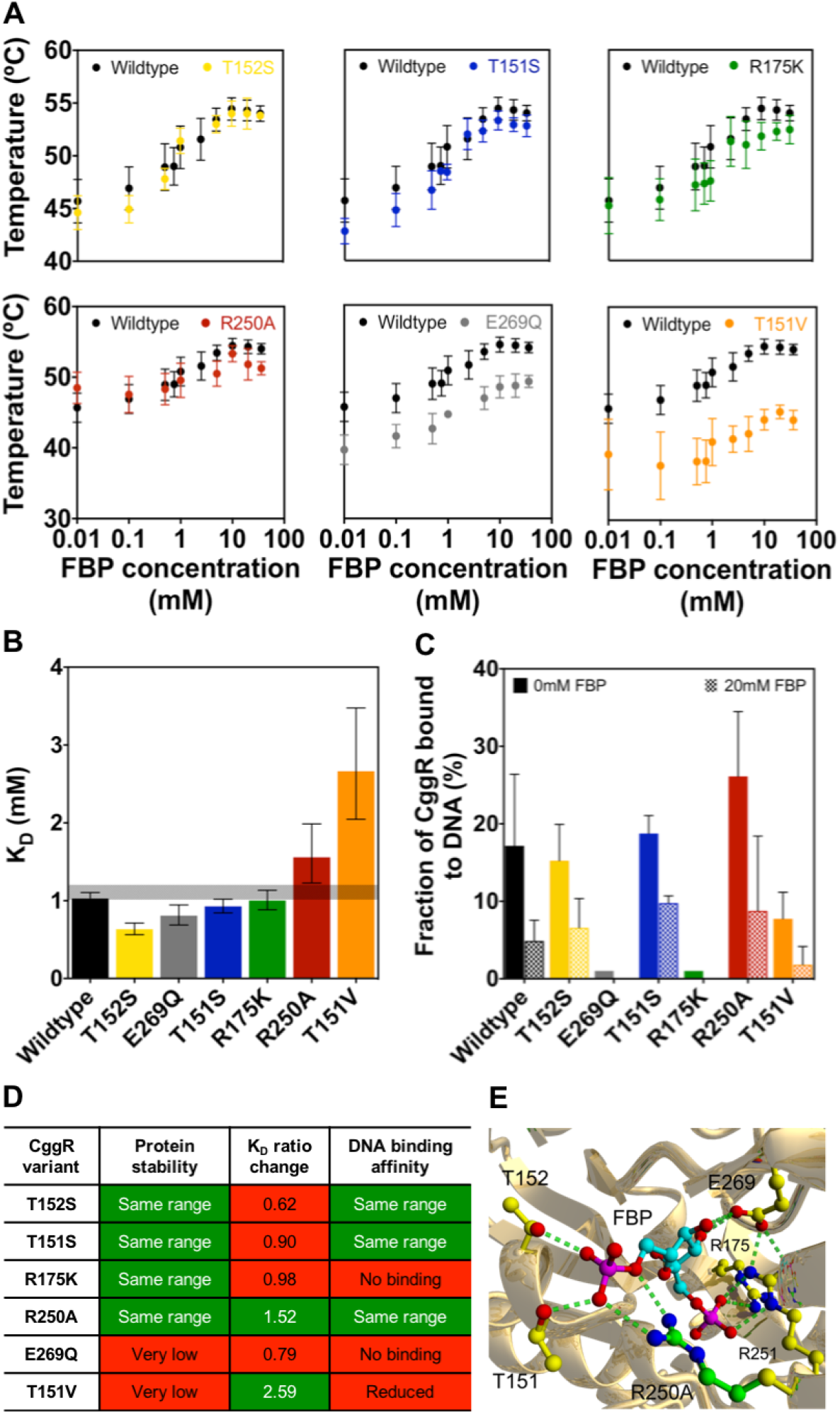
Biochemical characterization of CggR and respective mutants. (A) Thermal shift assays were used to determine the melting curves of wild-type CggR and mutants. Error bars correspond to the standard deviation of at least 5 replicates. (B) CggR-FBP affinity constants (K_D_’s) of the wildtype and mutant variants, determined by fitting a simple cooperative binding model to the melting curves data. Error bars indicate the 95% confidence intervals. (C) Quantification of CggR binding to DNA. The CggR bound to DNA fraction was calculated by dividing the intensity of the protein-DNA complex band by the total DNA. The background-subtracted total intensities of the CggR-DNA complex and the free-DNA bands were assessed with ImageJ. Error bars correspond to the standard deviation of three replicate experiments for the mutants and six for the wildtype. (D) Summary of the biochemical characterization of the wildtype and mutant CggR variants. K_D_ ratio change indicates the ratio between the K_D_ of the CggR mutant variant and the one of the wildtype. The desired characteristics achieved in the mutants are highlighted in green; in red the undesired effects of the mutations. (E) Ray-traced picture of the wildtype CggR and R250A mutant structure. Carbon atoms of the FBP ligand are in turquoise while the part of the side chain that is eliminated by the R250A mutation is in green. Hydrogen bonds are indicated with dashed lines.

To assess the DNA-binding capacity of the generated mutants, we performed Electro Mobility Shift Assays. We first measured the percentage of CggR bound to DNA in the absence of FBP, reflecting CggR’s binding affinity to DNA. Here, we found that the mutants R175K and E269Q variants did not bind to the DNA anymore (Figure 5C), and the mutant T151V only bound with lower affinity. The other mutants had a comparable DNA binding as the wildtype CggR. Next, using high (i.e. saturating) FBP levels (20 mM) to maximally promote release of CggR from the DNA and comparing the ratio between the percentage of the CggR bound to DNA, obtained at 0 mM of FBP, divided by the bound fraction at 20 mM, we found that only the R250A variant behaved similarly to the wildtype, with around 30% of the CggR remained bound to DNA at high FBP levels (Supplementary Figure 1).

Thus, as the R250A mutant fulfilled all desired criteria (Figure 5D), i.e. showed the desired decrease in FBP affinity, had a similar stability and DNA binding capability as the wildtype CggR, we selected this mutant for the flux sensor. This mutant had the additional advantage that it showed a flattened sigmoidal binding curve (Figure 5A), which is ideal for a sensor that needs to respond to a broad (cf. Figure 2A) FBP concentration range. This mutant eliminates the guanidine side chain of R250, which forms two H-bonds with the 6-phosphate group of FBP (Figure 5E).

### Testing the glycolytic flux sensor

Next, we implemented the flux-sensor elements in the wildtype and mutant (TM6) strains using either the wildtype CggR or the R250A mutant, genomically integrated in the HO locus under the control of the TEF1 promoter mutant 7 (P_TEFmut7_)^47^. We added a centromeric plasmid with the cis-regulatory CggR element (CggRO) controlling YFP (eCitrine) expression. To normalize the YFP signals for extrinsic cell-to-cell variation in the global state of the gene and protein expression machinery, we added a second fluorescent reporter protein RFP (mCherry) to the plasmid, under the control of the constitutive P_TEFmut8_ promoter^47^. Through proteome analyses we confirmed that CggR and mCherry expression levels correlate (Supplementary Figure 2), validating the use of mCherry to normalize YFP expression.

To determine the sensor output, i.e. the CggR activity, we quantified the YFP and mCherry fluorescence levels by flow cytometry, in the two strains grown on the different carbon sources. Growth rate analyses demonstrated that expression of the sensor constructs did not alter growth (Supplementary Figure 3). The ratio between the YFP and mCherry fluorescence, each corrected for autofluorescence determined by FACS, yielded the CggR repressor activity. Consistent with our design concept and the expected FBP-dependent de-repression of the synthetic promoter, here, we found a strong positive correlation between the YFP/mCherry ratios and the intracellular FBP levels (Figure 6A) and glycolytic flux (Figure 6B). This correlation was absent in control strains lacking CggR (Figure 6A and B). Furthermore, consistent with the growth-rate independent design of the sensor to respond solely to FBP levels, we found no correlation between the sensor output and the growth rate (Supplementary Figure 4).

**Figure 6.**
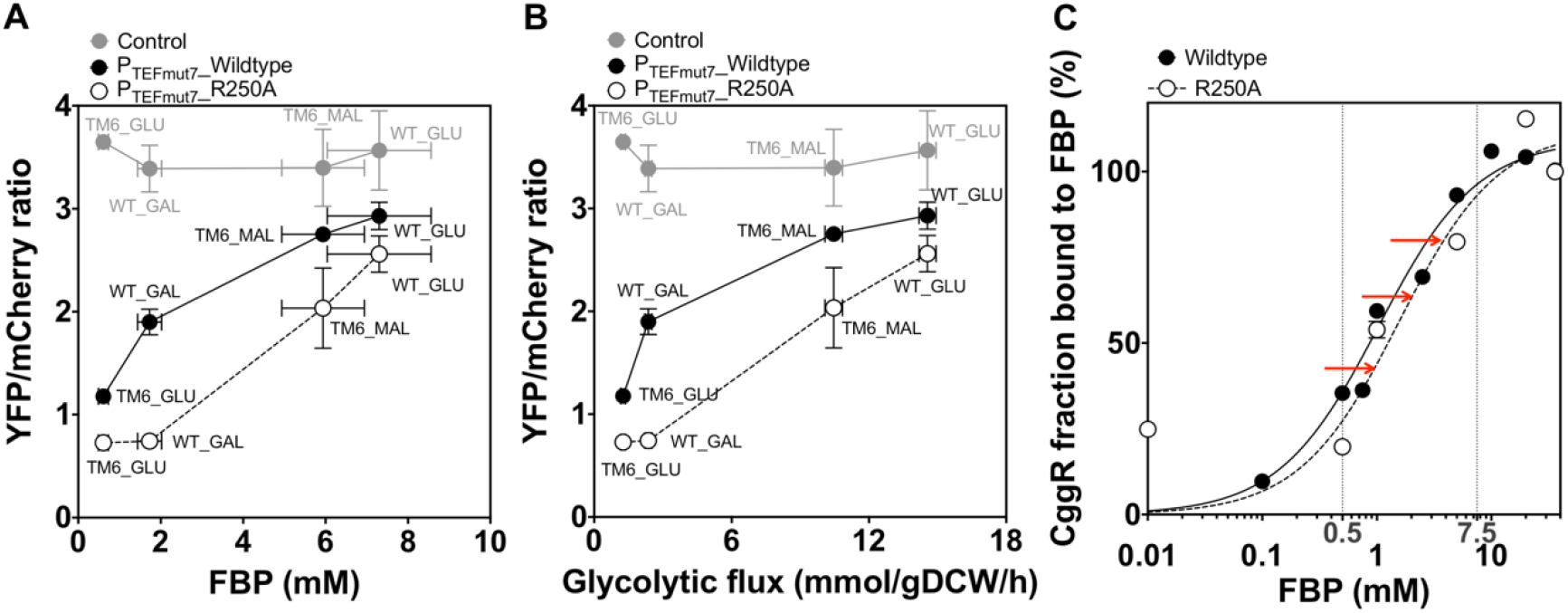
The engineered flux-sensor accurately reports glycolytic flux with a high dynamic flux range. Reporter activity of the sensor across (A) multiple FBP levels and (B) glycolytic fluxes. The glycolytic flux is here reported as the flux between the metabolites fructose 6-phosphate (F6P) and fructose-1,6-bisphosphate (FBP). Glycolytic fluxes were here estimated on the basis of physiological and metabolome data and a novel method to estimate intracellular fluxes^43^. Reporter activity is given by the YFP/mCherry ratio, calculated through the quantification of YFP and mCherry fluorescence along culture time using flow cytometry. Both YFP and mCherry fluorescence levels were first corrected for background using the same strains harboring the YCplac33 plasmid (Supplementary Table 3). The control is the wildtype and TM6 strains expressing only the reporter plasmid without CggR. Error bars represent the standard deviation of at least three replicate experiments. (C) Fraction of CggR bound to FBP across FBP concentrations. The red arrows indicate the shift in the percentage of CggR bound to FBP achieved in the R250A variant. The percentage of CggR molecules bound to FBP was calculated after normalizing the T_m_ values for unbound/bound state using the T_m_ at 0 mM FBP as unbound and at 36 mM (corresponding to maximum FBP concentration used) as total bound states. The curve fitting of the normalized values of CggR fraction bound to FBP was performed using a one-site specific binding model in GraphPad. The solid line corresponds to the wildtype CggR and the dashed line to the R250A variant. Vertical lines delimit the physiological FBP range.

While the sensor with the wildtype CggR (i.e. not optimized for FBP binding affinity) displayed also a correlation with FBP levels, the optimized version (in line with its lower FBP affinity) displayed a dynamic response that better covered the physiological concentration range of FBP (Figure 6A), and thus has a better capability to distinguish different glycolytic flux values (Figure 6A). When we estimated the wildtype and R250A CggR fraction bound to FBP, we observed that the main differences occurred at intermediate FBP levels (between 1.5 and 2.5 mM), in agreement with the fact that the K_D_ values of the two CggRs are around these FBP concentrations (Figure 6C).

It is remarkable that a single point mutation in CggR (R250A) led to such a different response curve (cf. Figure 6A). Consistent with this altered response curve, this point mutation solely altered the affinity to FBP (Figure 5B), but not its DNA-affinity (Figure 5C), stability (Figure 5D) nor its cellular abundance (Supplementary Figure 5). Thus, as only the affinity of FBP to CggR was altered, while everything else was kept constant, including the expression levels of CggR (Figure 4B), this demonstrates that the sensor’s output solely depends on the changing FBP. Therefore, together these data demonstrate that we have generated a sensor for FBP, and, as FBP levels correlate with glycolytic flux (Figure 2A), in fact a sensor that robustly and specifically reports glycolytic flux. While the wildtype and the mutant (TM6) strains used here have very different glycolytic flux levels during growth on glucose (Figure 2A), notably, our sensor unmasks this difference even though the environment was identical.

Altogether, this demonstrates that the recorded fluorescence ratio specifically responds to FBP levels. This means that we have generated a sensor that reports glycolytic flux.

### Proof of concept: quantification of glycolytic flux in coexisting quiescent and growing cells

Next, we aimed to test the engineered flux sensor with regards to its capability to detect single-cell differences in glycolytic flux when using microscopy, offering the possibility to co-assess other parameters, such as growth and cell division. First, we confirmed that also with the microscopic setup the output of the flux sensor still displays a linear correlation with glycolytic flux across conditions and strains (Figure 7A, Supplementary Figure 6A).

**Figure 7.**
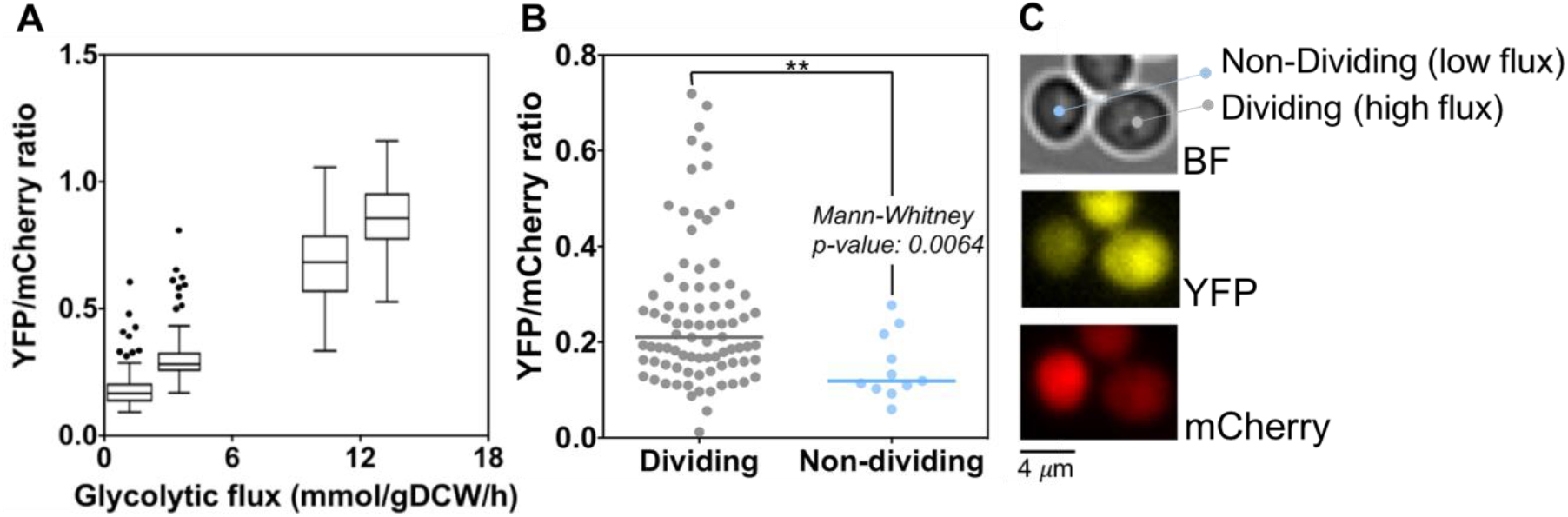
The glycolytic flux sensor can measure glycolytic flux in individual cells. (A) Tukey boxplots showing the YFP/mCherry ratio of individual cells measured by microscopy as a function of glycolytic flux. At least 35 cells were analyzed in each condition. The glycolytic flux is here reported as the flux between the metabolites fructose 6-phosphate (F6P) and FBP. Glycolytic fluxes were here estimated on the basis of physiological and metabolome data and a novel method to estimate intracellular fluxes^43^. (B) YFP/mCherry ratio measured by microscopy in co-existing dividing (high-flux) versus non-dividing (low-flux) isogenic TM6 cells on 10 gL^-1^ glucose. (C) Bright Field (BF), YFP and mCherry microscopy images for a co-existing dividing (high-flux) and a non-dividing (low-flux) TM6 cell expressing the flux sensor in 10g L^-1^ glucose minimal medium.

Next, we used microfluidics and time-lapse microscopy to cultivate the TM6 strain on glucose, where we recently showed that dividing cells with high flux co-exist with a small isogenic fraction of non-dividing cells with low glycolytic flux levels^50^. Here, using our new sensor, we found that non-dividing cells had indeed significantly lower YFP/mCherry ratios, even visibly by eye, compared to their coexisting dividing counterparts (Figure 7B and 7C, Supplementary Figure 6B), in line with their lower glycolytic flux. These results demonstrate that our flux sensor can be also used with microscopy, and is thus suitable for discrimination of individual cells with regards to their glycolytic flux levels even within clonal cell populations.

## Discussion

Here, exploiting the flux-signaling metabolite fructose-1,6-bisphosphate and the bacterial transcription factor CggR, we developed a biosensor, which allows to measure glycolytic flux in individual yeast cells. These engineering efforts, for which we used computational protein design, biochemical, proteome and metabolome analyses, entailed (i) development of a synthetic yeast promoter regulated by the bacterial transcriptional factor CggR, (ii) engineering of the transcription factors’ FBP binding site towards increasing the sensor’s dynamic range, and (iii) establishment of growth-independent CggR expression levels. Through single-cell time-lapse fluorescence microscopy experiments, we demonstrated that the sensor can reveal differences in glycolytic flux between single co-existing quiescent and growing cells.

Biosensor development based on transcription factors has recently seen rapid development^31, 51–53^. Three aspects of our work are worth to be highlighted: As no endogenous FBP-binding transcription factor is known in yeast, we had to transfer the *B. subtilis* transcription factor CggR into yeast. However, unlike previous studies which transplanted bacterial transcription factors into eukaryotes^20, 39, 54^, to ensure full orthogonality of the introduced sensing system to the host, we avoided the use of yeast-endogenous elements, such as DNA binding domains or chimeric fusions of TF with transcriptional activation domains, and the use of a nuclear localization sequences. Instead, in our design, we build the promoter from scratch, and exploited the natural mode of action of the TF also in the new host. Specifically, we used the ability of CggR to dimerize to allow for an effective repression mechanism also in yeast. Furthermore, and also in contrast to previous promoter engineering approaches, which often employed FACS-based screening approaches with large promoter libraries^19^, our approach was not a screening but a rational design approach. Our successful de novo engineering of a cis-regulatory element demonstrates that rational promoter development is possible when taking crucial factors into account, such as positioning and number of cis-regulatory elements, the transcription factor’s mode of action and the genomic context (for example, nucleosome positioning).

A second important aspect in our biosensor development was that we made sure that the output of the sensor is not influenced by growth-dependent changes in the transcription factor’s expression level, as its concentration also determines the synthesis rate of the gene product, and thus the output signal. In previous work, this point has mostly been ignored and TFs were typically ‘constitutively’ expressed, although constitutive expression does not necessarily lead to constant expression level across different growth rates^55^. Constant and condition-independent levels of the transcription factor is particularly important in the light of a glycolytic flux sensor, which likely will be applied across growth conditions. To accomplish growth rate-independent expression levels, using quantitative proteomics, we found that the P_TEFmut7_ promoter leads to condition-independent levels of CggR, while two other tested constitutive promoters, *i.e.* P_CMV_ and P_TEFmut2_, showed strong growth-dependent expression levels. We hope that future transcription-dependent biosensors will also consider the expression level of the TF as an important element in the development of the sensor.

Another important aspect in our biosensor development was the optimization of the biosensor’s dynamic range with regards to the sensed FBP levels and thus glycolytic fluxes. Here, we lowered the CggR-FBP binding affinity to cover the range of the intracellular FBP levels. Optimizing TF-effector sensitivity is not trivial, because transcription factors contain both an effector binding domain and a DNA-binding domain, which should not be altered when engineering the former. Here, we applied a semi-rational design approach, supported by computationally-guided protein design, to select mutants with lower FBP-CggR binding activity and unaltered CggR-DNA binding capacity. Demonstrating the challenge, only one mutation (R250A), out of a pool of 11 mutants, showed both desired features. Relevant for future engineering, only the mutations where the H-bond forming side chains were eliminated (R250A and T151V) resulted in the desired affinity loss. Furthermore, computational modeling with FoldX on the basis of available X-ray structures of both the normal and the activated conformation of the CggR effector-binding domain allowed to predict the instability or DNA-loss binding of some variants, indicating that computational stability predictions can successfully eliminate at least some dysfunctional mutants.

To construct the biosensor for glycolytic flux in yeast, we exploited the function of fructose-1,6-bisphosphate as a flux-signaling metabolite^30^ and we took advantage of the fact that FBP modulates the conformation of the *B. subtilis* transcription factor CggR^33^. Can a similar approach be pursued to develop flux sensors also for other metabolic pathways? This seems possible: On the basis of metabolite dynamics assessed across various nutrient conditions and known metabolite-protein and metabolite-RNA interactions, we recently compiled a list of several other candidates of flux-signaling metabolites^27^, which includes citrate, alpha-ketoglutarate, phosphoenolpyruvate, pyruvate and succinate. Exploiting respective metabolite-binding transcription factors, engineering biosensors for these metabolites should yield flux sensors also for other pathways. For instance, for citrate, whose concentration correlates with the cellular nitrogen flux^56^, the transcriptional activator CitI from lactic acid bacteria^57^ would be an excellent starting point. Thus, as flux-signaling metabolites also exist for other metabolic pathways^27^, and transcription factors exist for many of these metabolites^58^, it should be possible to develop flux biosensors also for other metabolic pathways.

We envision that our glycolytic flux biosensor, applicable in single yeast cells, will find numerous applications in fundamental research with *Saccharomyces cerevisiae*, i.e. to address the daunting emergence of metabolic heterogeneity and to in general investigate different metabolic phenotypes in a population, such as they occur during replicative aging^59^ or the cell cycle^3, 25^. Furthermore, we expect that the biosensor will also have strong value for applied research. Metabolic heterogeneity may be a significant problem in industrial fermentations, especially those with cell recycling as applied in beer brewing and bioethanol production^60–62^, where physiological and genetic changes can cause losses in fermentation performance^63^. In such large-scale yeast applications, our glycolytic flux sensor will provide a tool to study how and why metabolic sub-populations with high or low glycolytic flux phenotypes emerge. Beyond, we expect that the biosensor will find its application also as a screening tool in metabolic engineering efforts, for instance to screen for highly-productive phenotypes, rather than just selecting on growth.

## Materials and methods

### Generation and cloning of the CggR cis-regulatory element and reporter plasmid

The overall architecture of the synthetic promoter is outlined in the main text. The total DNA fragment size of the synthetic promoter element was 562 bp. This DNA fragment included two ends complementary to the reporter plasmid to allow for the Gibson assembly of the reporter plasmid and the synthetic promoter. The complementary flanking sites of the promoter element had a size of 100 bp at the 5’ end and 145 bp at the 3’ end. The initial sequence of the CggR cis-regulatory element was optimized using a computational method to minimize nucleosome affinity^40^. The functional elements of the CggR operator site, the TATA boxes and the 5’UTR were excluded from the sampling procedure. The algorithm was run for 75 rounds. The version 38 of the optimized CggR cis-regulatory element was selected since it showed the lowest affinity to nucleosomes and it was still feasible to be synthesized. DNA synthesis was performed with a STRING^TM^ DNA fragment (GenArt^TM^, ThermoFisher Scientific, MA, USA) and directly used for the assembly of the reporter plasmid. The functional DNA sequences of the CggR cis-regulatory element and their distance from the 5’ end of the synthesized fragment are listed in Supplementary Table 1.

To account for extrinsic cell-to-cell variation in the state of the gene and protein expression machinery, a constitutively expressed mCherry reporter was inserted into the low copy p416-loxP-KmR-TEFmut6-yECitrine centromeric plasmid^47^. The mCherry cassette included the mCherry ORF, the constitutive *TEF1* promoter mutant 8 (P_TEFmut8_), and the *ADH1* terminator, amplified from the plasmids pBS35, p416-loxP-KmR-TEFmut8-yECitrine and pWHE601 (Supplementary Table 3), respectively, with the primers listed in Supplementary Table 4. The three DNA fragments, *i.e.* P_TEFmut8_, mCherry and *ADH1* terminator, were assembled by PCR using the Phusion® High-Fidelity DNA Polymerase (New England Biolabs, MA, USA) and the mCherry_KpnI_fw and mCherry_KpnI_rv primers (Supplementary Table 4). The resulting 1.4 bp DNA fragment was purified, digested with KpnI and purified again with PCR Clean-up kit (Macherey-Nagel, Germany). The p416-loxP-KmR-TEFmut6-yECitrine plasmid was linearized with KpnI and dephosphorylated with Antarctic Phosphatase (New England Biolabs, MA, USA), and ligated with the mCherry expression cassette by T4 DNA ligase (New England Biolabs, MA, USA). The ligation assay was transformed into chemical competent *E. coli* DH5alpha cells. Clone screening was performed by sequencing the extracted plasmids with the primers listed in Supplementary Table 4.

To construct a plasmid carrying the regulated CggR promoter, we used the p416-loxP-KmR-TEFmut6-yECitrine with the inserted mCherry cassette (pTEF6-7) (Supplementary Table 3) and replaced the P_TEFmut6_ promoter by the CggR cis-regulatory element using Gibson assembly. The backbone of the pTEF6-7 plasmid was divided into three fragments, which were amplified by PCR using the primers listed in Supplementary Table 4. The three backbone fragments were combined together with the synthesized CggR cis-regulatory element using the NEB Gibson assembly kit (New England Biolabs, MA, US) according to the manufacturer instructions. 5 µl of reaction mix was transformed into chemical competent *E. coli* cells. Clone screening was performed to isolate the correct assembled pCggRO-reporter plasmid. The plasmid sequence was verified by Sanger sequencing of the extracted plasmids with the primers listed in Supplementary Table 4.

### Cloning of the CggR regulator and its variants and promoters

The open reading frame of the transcription regulator CggR of *B. subtilis* was codon optimized for expression in *S. cerevisiae*. A His_6_ and an Xpress epitope tag were added at the N-terminus of the protein. The CggR sequence was assembled from synthetic oligonucleotides (Thermo Fisher Scientific GeneArt AG, Germany) and the ORF was sub-cloned in the pET100/D-TOPO express cloning vector. Next, a Gibson assembly (New England Biolabs, MA, USA) was carried out to generate an expression cassette of the *cggR* gene for further integration in the HO-locus of *S. cerevisiae* genome. To allow for integration, the *cggR* gene was cloned into the integrative plasmid HO-poly-KanMX4-HO, where the KanMX4 resistance marker was replaced by the *ble* resistance gene.

Promoter and terminator of the *cmv* gene were amplified from the pCMV149 with a 5’ and 3’ overhang of a homologous sequence to the codon optimized *cggR* gene. The three DNA fragments, *i.e.* the *cggR* gene, the CMV promoter (P_CMV_) and terminator, were combined together by PCR using the Phusion® High-Fidelity DNA Polymerase (New England Biolabs, MA, USA). The resulting fragment was 2.2 kbp was gel-purified. A Gibson assembly was carried out to link all DNA fragments and assemble the final integrative pHO_pCMV_CggR_ble plasmid using the NEB Gibson assembly kit (New England Biolabs, MA, US). 5 µl of reaction mix was transformed into chemical competent *E.coli* DH5alpha cells. Clone screening was performed by sequencing the extracted plasmids. All the plasmids and primers used to generate the integrative plasmids are listed in Supplementary Tables 3 and 5, respectively. Additionally, the constitutive promoters TEF mutant 2 (P_TEFmut2_) and mutant 7 (P_TEFmut7_)^47^ were also cloned for testing the effect of different expression CggR levels on the biosensor output. P_TEFmut2_ and P_TEFmut7_ were amplified from p416-loxP-KmR-TEFmut2-yECitrine and p416-loxP-KmR-TEFmut7-yECitrine, respectively, and used to replace the P_CMV_ in the pHO_pCMV_CggR_ble plasmid.

To address the effect of a lower K_D_ towards FBP, the CggR mutant variant R250A was inserted in the above generated plasmids (replacing the CggR wt) using the NEB Gibson assembly kit (New England Biolabs, MA, US). 5 µl of reaction mix was transformed into chemical competent *E. coli* cells. Clone screening was performed by sequencing the extracted plasmids. The set of primers used for each integrative plasmid generation is listed in Supplementary Table 5. The strains generated and used throughout this work are listed in Supplementary Table 6.

### Cultivation and experimental sampling

All strains were cultivated in 500 mL Erlenmeyer shake-flask containing 50 mL of minimal medium^64^ inoculated with exponentially growing yeast cells to an initial OD_600_ of 0.1-0.2 (ca. 1-2×10^7^ cells). To adapt to the carbon source and to ensure metabolic steady state, two pre-culturing steps were carried out prior to the main culture. The inoculum was prepared in the identical minimal medium. All cultivations were performed at 30 °C and cultures were continuously shaken at 300 rpm. The medium was buffered at pH 5 with 10 mM KH-phthalate. Cells were cultured in different carbon sources that would generate distinct glycolytic fluxes and, as a consequence, FBP levels. Specifically, WT cells were grown in minimal medium containing 10 gL^-1^ of glucose, galactose or pyruvate and TM6 cells were cultured in minimal medium containing 10 gL^-1^ of maltose, glucose or pyruvate.

Cell counts were performed by flow cytometry (BD Accuri™ C6 flow cytometer, BD Biosciences, CA, USA) every hour for glucose, galactose and maltose. YFP and mCherry expression was assessed by measuring the fluorescence along culture time using flow cytometry through the FL1-A and FL3-A filters, respectively. Autofluorescence was assessed by measuring the fluorescence in cells containing the centromeric yeast plasmid YCplac33^65^ as a control using the FL1 and FL3 filters.

To determine production and consumption rates of metabolites during the cultivation, supernatant samples were taken every hour from the cultivations. 0.3 mL of the broth sample were centrifuged at 13 rpm for 2 min to separate the cells from the supernatant. The supernatant was transferred to filter columns (SpinX, pore size 0.22 µm), spun shortly, and stored at −20 °C until HPLC analyses were performed. At the end, the yeast dry mass was determined by filtering a certain volume of culture through pre-weighed nitrocellulose filters with a pore size of 0.2 µm. Filters were washed once with distilled water and kept at 80 °C for two days. Afterwards, they were weighed again. The cell dry mass at every measurement point was re-calculated from cell count and the dry mass cell count/ratio using the dry mass cell count/ratio obtained at the end of the fermentation.

### Quantification of physiological parameters

Glucose, pyruvate, glycerol, acetate and ethanol concentrations in the cultivation supernatant were determined by HPLC (Agilent, 1290 LC HPLC system) using a Hi-Plex H column and 5 mM H_2_SO_4_ as eluent at a constant flow rate of 0.6 mL min-1. The column temperature was kept constant at 60 °C. A volume of 10 µL of standards and samples was injected for analysis. Substrate concentrations were detected with refractive index and UV (constant wave length of 210 nm) detection. The chromatogram integration was done with Agilent Open Lab CDS software. Substrate and metabolite concentration were calibrated prior to the analysis of the fermentation samples using HPLC standards, which included all metabolites, relevant for the various conditions. The external standards covered the metabolites’ concentration range which was observed from the start until the end of the fermentation.

Carbon uptake rate calculations were performed using the time course data of the exponentially growing cultures of the different strains and carbon sources. From at least three independent biological replicates, the extracellular rates were estimated from measured concentration time-courses, e.g. glucose, ethanol, acetate, glycerol, pyruvate, and biomass, of the batch cultivation. Extracellular rates were estimated by fitting the concentration time courses to a mathematical model assuming exponential growth and constant yields in the culture. The regression and parameter estimation were implemented in gPROMS Model Builder v.4.0 (Process Systems Enterprise Ltd.).

### Quantification of intracellular metabolite levels

A sample of 3×10^7^ cells was taken and immediately quenched in 10 mL methanol, which was beforehand cooled down to −40°C. The cells were separated from the organic solvent by centrifugation (5 min, 21000 g, 4 °C), washed with 2 mL methanol, separated again by centrifugation and stored at −80 °C. For the following analysis the cell pellet was re-suspendend in extraction buffer (methanol, acetonitrile and water, 4:4:2 v/v/v supplemented with 0.1 M formic acid) and an internal standard of ^13^C-labeled metabolites was added to the extraction. This standard was obtained and quantified from exponentially grown cell cultures prior to the experiment. The extraction was agitated for 10 min at room temperature and thereafter centrifuged at maximum speed. The supernatant was transferred to a new vial and the cell pellet re-suspended in extraction buffer and the extraction procedure was repeated a second time. The supernatants from both steps were combined and centrifuged for 45 min at 4 °C and 21000 g to remove any remaining non-soluble parts. Thereafter, the supernatant was vacuum dried at 45 °C for approximately 1.5 h and subsequently re-dissolved in 200 µL water.

The extracted intracellular metabolites were identified and quantified using a UHPLC-MS/MS system as done previously^66^. Specifically, the chromatographic separation was performed on a Dionex Ultimate 3000 RS UHPLC (Dionex, Germering, Germany) equipped with a Waters Acquity UPLC HSS T3 ion pair column with precolumn (dimensions: 150 x 2.1 mm, particle size: 3 μm; Waters, Milford, MA, USA). The injection volume was 10 μL and the samples were permanently cooled at 4 °C. A binary solvent gradient was employed (0 min: 100%A; 5 min: 100%A 10 min: 98%A; 11 min: 91%A; 16 min: 91%A; 18 min: 75%A, 22 min: 75%A; 22 min: 0%A; 26 min: 0%A; 26 min: 100%A; 30 min: 100%A) at a flow rate of 0.35 mL min^-1^, where solvent A was composed of 5% (v/v) methanol in water supplemented with 10 mM tributylamine, 15 mM acetic acid and 1 mM 3,5-heptanedione and isopropanole as solvent B. The detection was done using multiple reaction monitoring (MRM) on a MDS Sciex API365 tandem mass spectrometer upgraded to EP10+ (Ionics, Bolton, Ontario, Canada) and equipped with a Turbo-Ion spray source (MDS Sciex, Nieuwerkerk aan den Ijssel, Netherlands) with the following source parameter: NEB (nebulizing gas, N2): 12 a.u., CUR (curtain gas, N2): 12 a.u., CAD (collision activated dissociation gas): 4 a.u., IS (ion spray voltage): −4,500 V, TEM (temperature): 500 °C.

The amounts of metabolites determined in each sample, were converted into intracellular concentrations, using the determined number of cells in the sample and the respective cell volume. The cell volume was determined by taking an image of mid-exponential cells of wildtype (WT) and TM6 in the various conditions. The cells were placed on a microscopic slide. Approximately 200 cells per condition and replicate were evaluated on several positions / images of the microscopic slide. The cell volume was estimated using budJ plugin of ImageJ^67^.

### Determination of intracellular fluxes

To estimate the glycolytic flux for the two strains and the different substrate conditions, we used a thermodynamic constraint-based metabolic model, and a new approach for metabolic flux analysis^43^. The model and constraints were based on what we previously used, but the network was extended by reactions describing the uptake and metabolization of galactose and maltose (Supplementary Table 8 and 9). The model was fitted to experimental data, which here comprised of intracellular metabolite concentrations, extracellular fluxes (as measured for the different conditions and strain) and standard Gibbs free energies of reaction (Δ_r_G°), determined from the component contribution method^68^. The experimental data of all six conditions are given in Supplementary Tables 10 and 11. The fitting was done as previously, i.e. jointly for all conditions to identify one condition-dependent set of Δ_r_G⁰, but without regularization. The result of the regression analysis and the goodness of fit are presented in Supplementary Figure 7. Next, we performed a flux variability analysis to determine the limits of the solution space of each flux for all growth conditions and the two strains, for which the fitted values of the measured extracellular fluxes were allowed to vary ±2.95 standard deviations. Within those limits, 1000 flux distributions were sampled for each condition with optGpSampler^69^ using linear approximations for the non-linear thermodynamic constraints, as we did previously^43^. The mean and standard error of each metabolic flux at each condition was determined from this sample population.

### Quantification of CggR and mCherry levels

Intracellular CggR and mCherry levels were quantified in CMV, P_TEFmut2_ and P_TEFmut7__CggR yeast strains. WT and TM6 cells were grown using the same scheme as described above. Cell counts were monitored by flow cytometry (BD Accuri™ C6 flow cytometer (BD Biosciences, CA, USA) and cells were harvested once they reached a concentration of 10^7^ cells mL^-1^. To do so, 10 mL of cells were pelleted (ca. 10^8^ cells per sample), washed twice with ice cold PBS (0.1 mM PO_4_^3−^, 1.37 mM NaCl, 0.027 mM KCl) and centrifuged at 10000xg, at 4 °C for 10 min. The cell pellet was frozen in liquid nitrogen and stored at −80 °C until further analysis. For each strain, three biological replicates and three technical replicates were taken.

The cell pellet was reconstituted in 40 µL 2% sodium-deoxycholate (SDC); 10 mM TCEP; 100 mM ammonium bicarbonate and sonicated two times for 10 seconds using a UP200St with VialTweeter (Hielscher Ultrasonics GmbH, Germany). Heat treatment was performed for 10 minutes at 95°C. After cooling, the protein concentration was determined by BCA assay for each sample (Thermo Fisher Scientific, MA, USA). Sample aliquots containing 100 µg protein were used for the following steps. Alkylation was performed by adding iodoacetamide to a final concentration of 15 mM and incubation for 45 minutes at RT, in the dark. The samples were diluted to 1% SDC using 100mM ammonium bicarbonate and mass spectrometry grade Trypsin (Promega, WI, USA) was added at a ratio of 1:50 (µg Trypsin:µg Protein) and samples were incubated overnight at 37 °C, 400 rpm. The reaction was stopped by adding trifluoroacetic acid to a final concentration of 1%. Sample cleanup by solid phase extraction was performed with Pierce® C18 tips (Thermo Fisher Scientific, MA, USA) according to the manufacturer instructions. The eluate fraction was dried under vacuum and reconstituted with 20 µL of a mixture of 2% acetonitrile and 0.1% formic acid.

1 µg of peptides of each sample were subjected to LC–MS analysis using a dual pressure LTQ-Orbitrap Elite mass spectrometer connected to an electrospray ion source (Thermo Fisher Scientific, MA, USA) as described recently^70^ with a few modifications. In brief, peptide separation was carried out using an EASY nLC-1000 system (Thermo Fisher Scientific, MA, USA) equipped with a RP-HPLC column (75 μm × 30 cm) packed in-house with C18 resin (ReproSil-Pur C18–AQ, 1.9 μm resin; Dr. Maisch GmbH, Ammerbuch-Entringen, Germany) using a linear gradient from 95% solvent A (0.15% formic acid, 2% acetonitrile) and 5% solvent B (98% acetonitrile, 0.15% formic acid) to 28% solvent B over 100 min and to 45% B over 20 min at a flow rate of 0.2 μl/min. The data acquisition mode was set to obtain one high resolution MS scan in the FT part of the mass spectrometer at a resolution of 120,000 full width at half-maximum (at m/z 400) followed by MS/MS scans in the linear ion trap of the 20 most intense ions using rapid scan speed. The charged state screening modus was enabled to exclude unassigned and singly charged ions and the dynamic exclusion duration was set to 60 s. The ion accumulation time was set to 300 ms (MS) and 25 ms (MS/MS).

For label-free quantification, the generated raw files were imported into the Progenesis QI software (Nonlinear Dynamics (Waters), Version 2.0) and analyzed using the default parameter settings. MS/MS-data were exported directly from Progenesis QI in mgf format and searched against a decoy database the forward and reverse sequences of the predicted proteome from *Saccharomyces cerevisiae* (strain ATCC 204508 / S288c, UniProt, download date: 15/12/2016) including common contaminants, like keratins) and CggR from *Bacillus subtilis* (strain 168, total of 14,248 entries) using MASCOT (version 2.4.1). The search criteria were set as follows: full tryptic specificity was required (cleavage after lysine or arginine residues); 3 missed cleavages were allowed; carbamidomethylation (C) was set as fixed modification; oxidation (M) as variable modification. The mass tolerance was set to 10 ppm for precursor ions and 0.6 Da for fragment ions. Results from the database search were imported into Progenesis QI and the final peptide measurement list containing the peak areas of all identified peptides, respectively, was exported. This list was further processed and statically analyzed using our in-house developed SafeQuant R script^70^. The peptide and protein false discovery rate (FDR) were set to 1% using the number of reverse hits in the dataset.

The CggR intracellular abundance is the cell protein content of CggR calculated using the iBAQ approach^71^. The relative abundance of CggR was calculated by normalizing the cell protein content of CggR to the cell protein content of the CggR measured on glucose and using same promoter.

### *In silico* design of a CggR mutant library with altered affinity for FBP

FoldX was used to predict the stability and structure of CggR mutants^49^. FoldX evaluates the change in folding energy (ΔΔG^fold^) due to a point mutation. This ΔΔG^fold^ equals the Gibbs folding energy (ΔG^fold^) of the mutant minus the Gibbs folding energy of the wild-type protein. The FoldX settings were as default for the calculation of the stability effects of point mutations, with averaging of five independent predictions per variant^72^. In the dimeric CggR structure (PDB id: 3BXF), two conformations occur because the binding of FBP to only one of the subunits causes a local conformational change^48^. To model the effect of mutations on the equilibrium between those two conformations, two dimer structures were generated, in which the same conformation occurs in both subunits. This was done by duplicating one subunit, superimposing the resulting copy on the coordinates of the other subunit by a least-squares fit, and then eliminating the original subunit at that position. Suitable residues to mutate in the FBP binding site were identified by visual inspection using Yasara^73^. From a total of 18 mutations predicted, 6 were selected to be tested *in vitro* (Table 1) based on the desired mild impact on the dissociation constant (K_d_) of CggR.

### Site-directed mutagenesis of CggR

For the generation of the CggR mutants with decreased FBP binding affinity, a site-directed mutagenesis approach based on PCR was performed. The set of primers containing the mutation of interest are listed in Supplementary Table 7. All the PCR reactions were performed according to the Phusion® High-Fidelity DNA Polymerase (New England Biolabs, MA, USA) protocol using 0.05 Units of Phusion and 100 µM dNTPs in a total volume of 25 µl.

To eliminate contamination of the original template DNA, a DpnI digestion was performed by adding 1 µl of DpnI to the PCR mix, followed by an overnight incubation at 37 °C. 9 µl of the DpnI digestion product was then transformed into chemically competent *E. coli* DH5alpha cells. Plasmid DNA extraction was done with the Nucleospin Plasmid purification kit (Macherey-Nagel, Germany). The confirmation of the mutated CggR variants was performed by sequencing of the extracted plasmids.

### CggR protein expression and purification

CggR wildtype and the generated mutants were cloned in pET100/D-TOPO (Thermo Fisher Scientific; Waltham, MA), with an N-terminal His_6_-tag and expressed in *E. coli*. All constructs were verified by sequencing. For protein production, a single colony was used to inoculate 50 mL LB containing 100 µg mL^-1^ ampicillin, and the culture was grown at 37°C overnight. This culture was diluted to an optical density (OD_600_) of 0.05 in a final volume of 2 L. Protein expression was induced at OD_600_ 0.5 by addition of 10 µM IPTG, and cells were kept at 30 °C and 180 rpm for four more hours. Cells were harvested by centrifugation at 6,675xg at 4 °C for 20 minutes and washed once with 30 mL of 50 mM Tris-HCL buffer (pH 7.2). Cells were pelleted at 3000xg at 4 °C for 40 minutes, frozen in liquid nitrogen and stored at −80 °C until further use.

For protein purification, cell pellets were thawed on ice and resuspended with 10 mL of ice cold lysis buffer (50 mM KH_2_PO4, 300 mM NaCl, 1 mM EDTA, pH 7.5) per gram of cell pellet, and lysed by high-pressure disruption (Constant Cell Disruption System, Ltd, UK) in one passage at 25 Kpsi at 4 °C. Prior to lysate centrifugation, 1 mM of PMSF, 20 mM MgCl_2_ and 10 µg/mL of DNase (Sigma-Aldrich, MO, USA) was added. Cell debris were removed by centrifugation at 35200xg at 4 °C for 20 minutes. The cleared lysate was incubated with 0.5 mL of a nickel sepharose resin (GE healthcare, Little Chalfont, UK), pre-equilibrated with 50 mM of KH_2_PO_4_ buffer (pH 7.5), and incubated at 4 °C in batch mode overnight. The nickel sepharose-lysate suspension was poured onto a 10 mL disposable column (Bio-Rad), and the settled resin washed with 20 column volumes of a first wash solution (50 mM KH_2_PO_4_, 300 mM NaCl, 60 mM imidazole, pH 7.5), followed by 20 column volumes of a second wash solution (50 mM KH_2_PO_4_, 300 mM NaCl, 50 mM L-histidine, pH 7.5). Protein elution was performed with 9 mL of elution buffer (50 mM KH_2_PO_4_, 300 mM NaCl, 235 mM L-histidine, pH 7.5).

Protein concentration was measured by absorbance at 280 nm and extinction coefficient of 4.2. Protein purity and integrity were evaluated by running the samples in a 10% SDS PAGE gel using protein concentrations of 0.1 mg/mL. A buffer exchange (50 mM KH_2_PO_4_, 300 mM NaCl, pH 7.5) was performed on the fractions with concentrations above 1 mg/mL and purity above 95%. The protein purified stocks were stored at 4 °C until needed.

### Thermal shift assays

A sample mixture of 25 µL final volume containing 5 µL of 5x SYPRO Orange (Molecular Probes; Eugene, OR), 0.2 mg of the purified CggR, and different concentrations of the FBP metabolite (0; 0.01; 0.1; 0.5; 0.75; 1; 2.5; 10; 20 and 36 mM) was prepared on ice. Control experiments were performed to test the effect of the counter ion NaCl present in FBP salt solutions. Here, the NaCl was added instead of FBP to a final concentration 3-fold higher than the FBP itself (0; 0.03; 0.3; 1.5; 2.25; 3; 7.5; 30; 60 and 108 mM).

Sample mixtures were transferred into 96-well PCR plates (Bio-Rad, CA, USA), sealed with Optical-Quality Sealing Tape (Bio-Rad, CA, USA), and analysed in a CFX96 Real-Time System combined with C1000 Touch Thermal Cycler (Bio-Rad, CA, USA). Analysis consisted of a single heating cycle from 20°C to 99°C with increments of 0.5°C steps, followed by fluorescence intensity monitoring with a charge-coupled device camera. The wavelengths for excitation and emission were 490 and 575 nm, respectively. The melting temperature (T_m_) was automatically calculated by the control software and corresponded to the local maximum of the first derivative of measured fluorescence versus temperature.

The K_D_ of the wildtype and mutant CggR variants was calculated by fitting the T_m_ data into a simple cooperative model using the GraphPad software.

### Electrophoresis mobility shift assay

Fluorescently-labeled DNA fragments were generated by hybridization of single-stranded forward (labeled with Alexa Fluor-647 at the 5’ end) and reverse (unlabeled) oligonucleotides containing the CggR operator sites. The hybridization protocol and respective hybridization efficiency control were performed as described elswhere^32^.

Hybridized labeled DNA fragments (final concentration 35 nM) were incubated 20 minutes at room temperature with 2 µM or 4 µM of the purified wildtype or mutant CggR variants, in the presence of different concentrations of FBP (0; 0.5; 1; 2.5; 5; 10; and 20 mM) in a final volume of 25 µL in binding buffer (10 mM Na_3_PO_4_ pH 7.8, 100 mM NaCl, 1 mM EDTA, 1 mM DTT, 5% glycerol) including 1 µg of salmon sperm DNA. The sample mixtures were loaded in a 5% native polyacrylamide gel and run in native conditions in the dark with constant voltage (200V) at 4 °C for 2 hours.

Fluorescence was imaged using the Typhoon 9400 (Amersham Biosciences, UK) with the excitation set to 650 nm and the emission set to 655 nm. The background-subtracted total intensities of the protein-DNA complex and the free-DNA bands were assessed using imageJ^74^, and bound DNA/total DNA was calculated by dividing the intensity of the protein-DNA complex band by the total DNA (i.e. the sum of the signal from the protein-DNA complex band plus the one from free-DNA).

### Time-lapse microscopic analyses

For testing the glycolytic flux-sensor output with microscopy, a microfluidic device^75, 76^ was used, which was loaded with cells at log phase growing in glucose, maltose or galactose. Monitoring of cells took place using an inverted fluorescence microscope (Eclipse Ti-E; Nikon). During the experiment, a custom-made microscope incubator (Life Imaging Services GmbH) retained the temperature constant at 30 °C, and cells were continuously supplied with fresh medium. For illumination, an LED-based excitation system (pE2; CoolLED) was used, and images were recorded with an Andor 897 Ultra EX2 EM-CCD camera using a CFI Plan Apo VC 60x Oil (NA = 1.4; Nikon) objective. 300 ms exposure time and 50% light intensity were used for YFP (500 nm excitation using a 520/20 nm excitation filter and a 515 nm beam-splitter, 535/30 nm emission filter, EM gain 25), and 200 ms exposure time and 25% light intensity were used for mCherry measurements (565 nm excitation using a 562/40 nm excitation filter and a 593 nm beam-splitter, 624/40 nm emission filter, EM gain 25). Brightfield images were recorded for cell segmentation.

For comparing the flux sensor output between coexisting high flux (dividing) and low flux (non-dividing) cells, TM6 cells from log-phase (10 gL^-1^ glucose) cultures were loaded in the microfluidic device, and after one initial round of fluorescence imaging, cells were followed only with the brightfield channel for 20 hours to determine budding activity.

Cell segmentation to determine mean cell YFP and mCherry fluorescence intensities was performed with the ImageJ plugin BudJ^77^ using the brightfield images. Before computing the YFP to mCherry ratio for each cell, fluorescent intensity measurements were corrected for background fluorescence by subtracting the modal grey value of the whole image area in each fluorescent channel. Statistical analyses and plotting were performed in GraphPad.

## Acknowledgements

The research of MH, GH, and FM has received funding from the European Commission under grant agreement no. 613745, PROMYS, and the one of MH and JS under grant agreement no. 675585, SymBioSys. Further, the authors would like to thank Silke Bonsing-Vedelaar for support during the metabolomics experiments, Marta Palka and Ivan Frak for their support during early measurement campaigns and cloning, Brenda Bley-Folly and Alvaro Ortega for their advice for on the implementation of the EMSA and thermal shift experiments, and Johan Hekelaar for preparation of the proteome samples.

## Author contributions

GH and MH conceived the idea of the study. FM, GH and MH designed the study. FM, GH, JN and AL performed the experiments. HJW performed the protein modeling analysis. JH and AS performed the proteomic analysis. JS performed the computational metabolic flux analysis, FM, GH, JN and AL analyzed experimental data. FM, GH, AL and MH wrote the manuscript. MH supervised the study.

## Data availability

All mass spectrometry raw data files have been deposited to the ProteomeXchange Consortium (accession code PXD012964, http://proteomecentral.proteomexchange.org, reviewer login: username, password: aVZoKObp) via the PRIDE partner repository^78^.

## Supplementary Information

**Supplementary Table 1.**
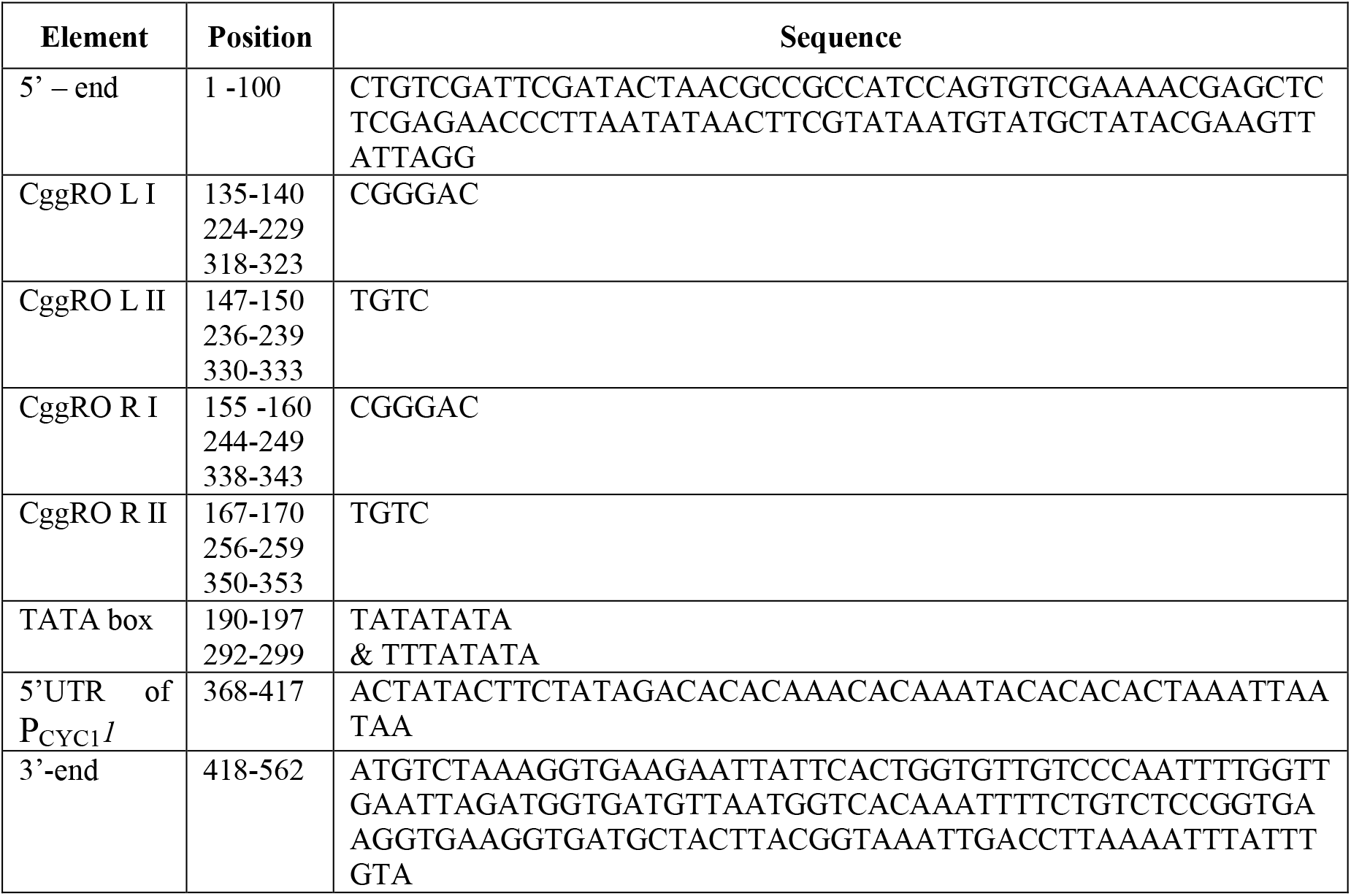
List of the position of the cis-regulatory element sequences.

**Supplementary Table 2.** Detailed motivations for the introduced mutations.

| Mutation | Expected effects of the mutations |
| --- | --- |
| **T151S** | T151 makes a hydrogen-bond via its hydroxyl hydrogen atom to a terminal oxygen of the 6-phosphate group of FBP. This H-bond appears to be strong, as evidenced by an oxygen-oxygen distance of 2.6 Å in the 3BXF structures. The T151S mutation could preserve this H-bond bond as the hydroxyl group of a serine can be positioned at the same spot as that of a threonine, although less rigidly due to the deletion of the methyl group of the threonine. Thus, this mutation is expected to have **negligible to mild** effects on binding affinity |
| **T151V** | This mutation introduces an isosteric replacement for the original threonine, and is thus expected to eliminate the above-mentioned strong H-bond and decrease binding affinity for FBP. One would therefore expect to see a **mild to strong** effect on the binding affinity for FBP. |
| **T152S** | Like T151, the hydroxyl side chain of T152 makes a relatively strong H-bond to one of the terminal oxygens of the 6-phosphate of FBP (oxygen-oxygen distance 2.6 Å). With the same rationale as for T151S, the effect of the mutation of FBP binding is expected to be **negligible to mild**. |
| **R175K** | The positively charged R175 side chain makes a strong (nitrogen-oxygen distance 2.7 Å) H-bond with a terminal oxygen of the 1-phosphate group of FBP. Mutating the arginine to a lysine may give less strong interactions to the FBP, resulting in decreased affinity. A complicating factor is that R175 radically changes side chain orientation upon binding of FBP due to a G177-Q185 loop movement in the protein. The FoldX predictions suggest that for the R175K mutant the equilibrium may shift to the FBP conformation also in the absence of FBP. Thus, the effect of the mutation could vary from **negligible to strong**. |
| **R250A** | R250 makes two H-bonds with the 1-phosphate group of FBP. The N-O distances are 3.3 and 2.9 Å. One H-bond is to a terminal phosphate oxygen, the other is to the ether oxygen (in between the phosphor and the carbon atom). One would expect the overall effect on binding to be strong unless the created cavity allows for water molecules to come in and replace the hydrogen bonds that were lost by removing the R250 side chain. Thus, mutation of this residue could give rise to a **mild to strong** affinity decrease. |
| **E269Q** | The side chain of E269 accepts H-bonds for the hydroxyl groups at the 3 and 4 positions of the FBP. The oxygen-oxygen distances are 2.6 and 2.5 Å, which indicates strong H-bonding interactions. Mutation to glutamine would possibly only change the H-bond pattern but not decrease the overall number of H-bonds. This would then still be expected to weaken the interactions as the newly formed H-bonds would be less polarized than those in the wild-type structure. Thus, the expected result of this mutation is a **mild to strong** affinity decrease for FBP |

**Supplementary Table 3.** List of the plasmids used and generated in this work.

| Plasmid | Function | Source |
| --- | --- | --- |
| pET100-CggR-Sc | pET100/D-TOPO plasmid used for CggR protein expression | This study |
| HO-poly-KanMX4-HO | Integrative plasmid | <sup>79</sup> |
| pUG66 | *E. coli* / *S. cerevisiae* shuttle vector containing, Amp<sup>+</sup>, loxP-bleR-loxP disruption cassette | <sup>80</sup> |
| pCM149 | Plasmid for integrative yeast transformation, marker LEU2, CMVp(tetR) | <sup>81</sup> |
| p416-loxP-KmR-TEFmut2-yECitrine | CEN/ARS plasmid, loxP-KanMX4-loxP, TEFmut2, yECitrine; Amp<sup>+</sup> | <sup>47</sup> |
| p416-loxP-KmR-TEFmut7-yECitrine | CEN/ARS plasmid, loxP-KanMX4-loxP, TEFmut7, yECitrine; Amp<sup>+</sup> | <sup>47</sup> |
| p416-loxP-KmR-TEFmut6-yECitrine | CEN/ARS plasmid, loxP-KanMX4-loxP, TEFmut6, yECitrine; Amp<sup>+</sup> | <sup>47</sup> |
| p416-loxP-KmR-TEFmut8-yECitrine | CEN/ARS plasmid, loxP-KanMX4-loxP, TEFmut8, yECitrine; Amp<sup>+</sup> | <sup>47</sup> |
| pHO_pCMV_CggR_ble | Plasmid for genomic integration of CggR and P<sub>CMV</sub> promoter | This study |
| pHO_pTEFmut2_CggR_ble | Plasmid for genomic integration of CggR and P<sub>TEFmut2</sub> promoter | This study |
| pHO_pTEFmut7_CggR_ble | Plasmid for genomic integration of CggR and P<sub>TEFmut7</sub> promoter | This study |
| pHO_pTEFmut2_CggR_R250A_ble | Plasmid for genomic integration of CggR and P<sub>TEFmut2</sub> promoter | This study |
| pHO_pTEFmut7_CggR_R250A_ble | Plasmid for genomic integration of CggR and P<sub>TEFmut7</sub> promoter | This study |
| pBS35 | Plasmid containing the mCherry ORF sequence | <sup>82</sup> |
| pWHE601 | Plasmid containing the ADH1 terminator sequence | <sup>83</sup> |
| pYCplac33 | Yeast centromeric plasmid for fluorescence background correction | <sup>65</sup> |
| pTEF6-7 | Constitutively regulated eCitrine and mCherry reporter plasmid | This study |
| pCggRO-reporter | Constitutively expressed mCherry and CggR-regulated YFP (eCitrine) reporter plasmid | This study |

**Supplementary Table 4.**
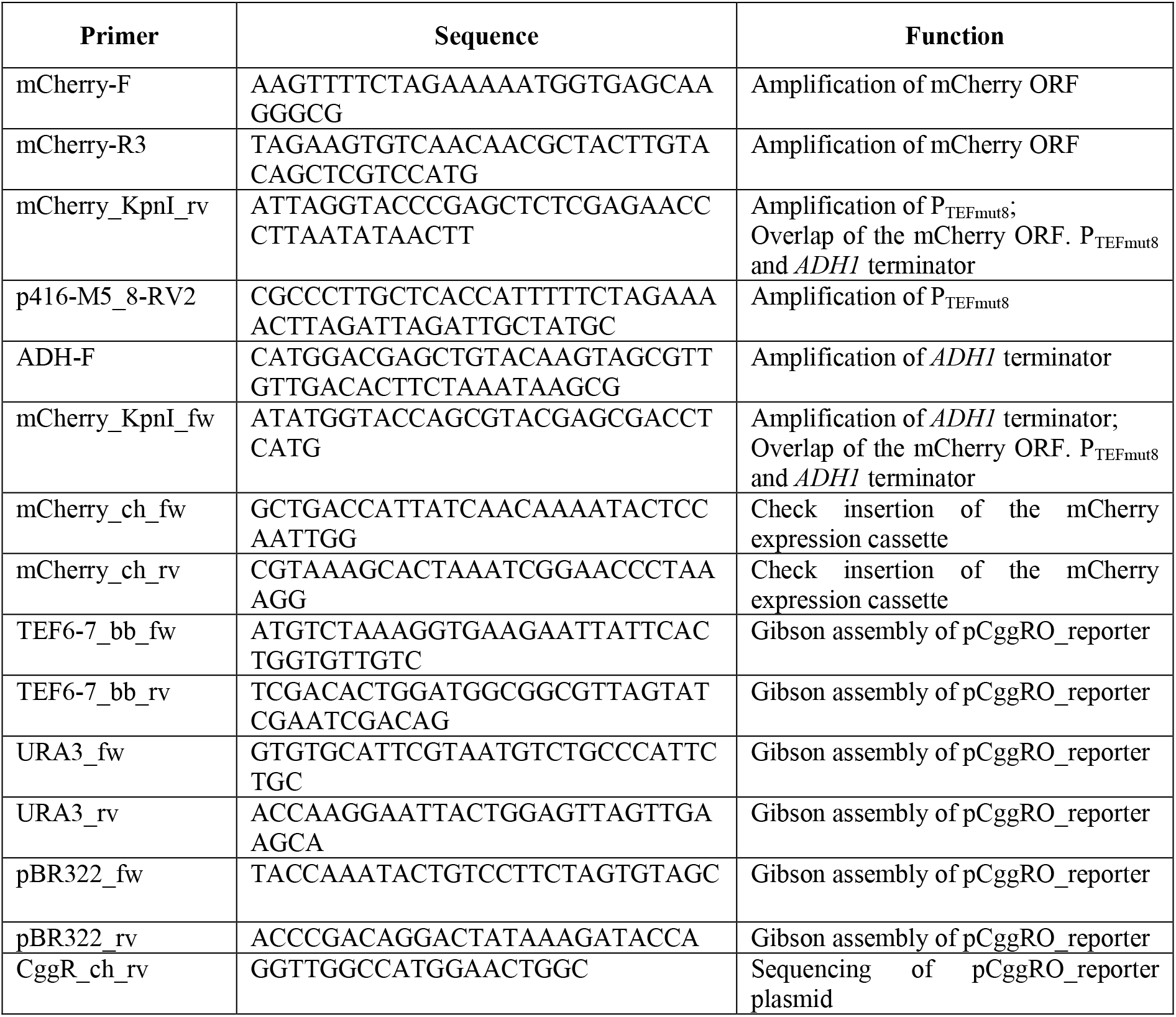
Primers list for the construction of the CggR cis-regulatory reporter plasmid.

**Supplementary Table 5.**
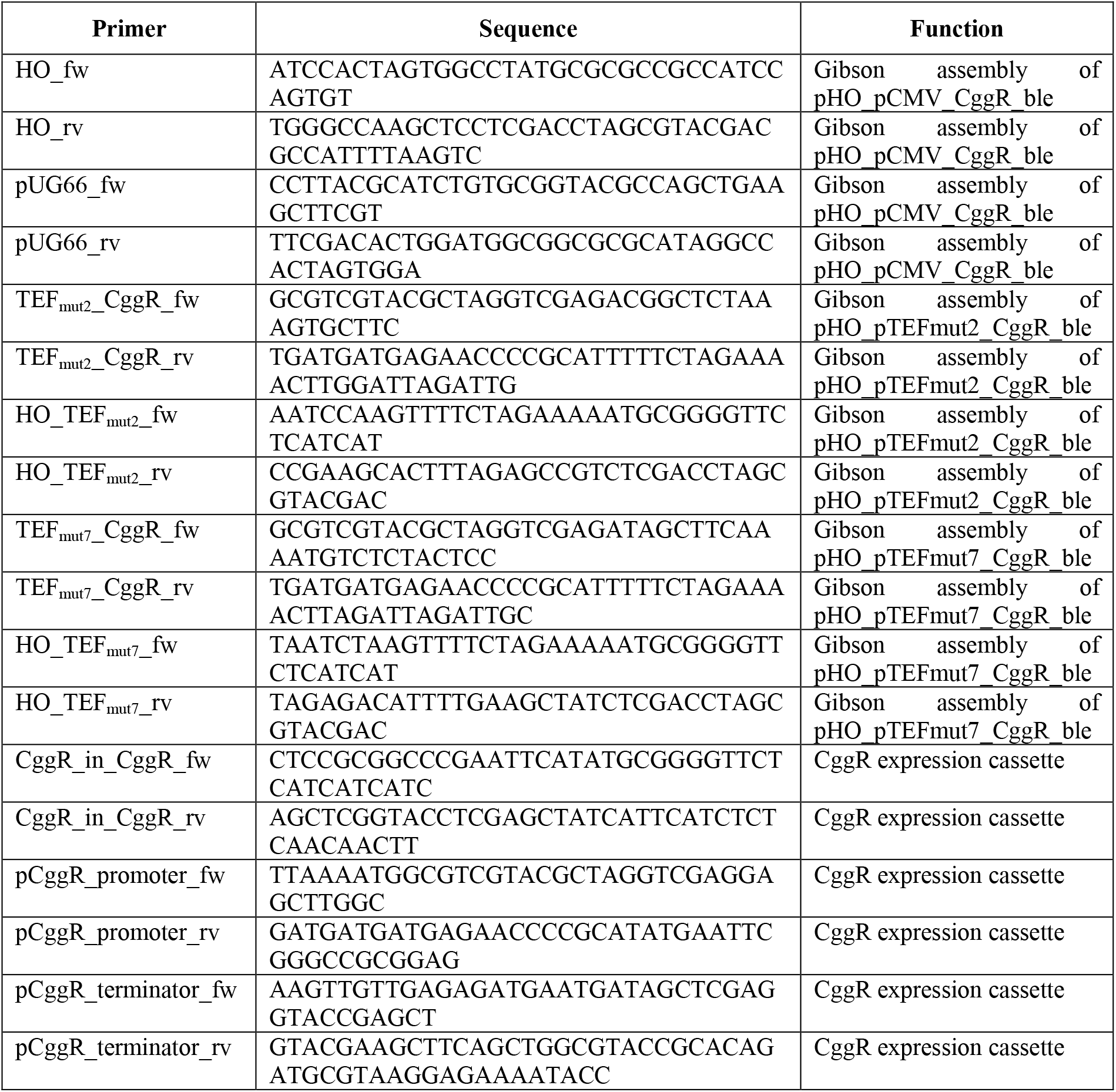
List of primers used to generate the CggR integrative plasmids.

**Supplementary Table 6.** List of yeast strains used and generated in this work.

| Strain name | Genetic alteration | Genomic insertion | Plasmid name | Function | Source |
| --- | --- | --- | --- | --- | --- |
| WT_pCggRO | Plasmid | *N.A.* | pCggRO | Wildtype strain containing the pCggRO reporter plasmid for control of the cis-regulatory dependent YFP/mCherry expression | This work |
| WT_YCplac33 | Plasmid | *N.A.* | pYCplac33 | Wildtype strain containing the YCplac33 plasmid for fluorescence background correction | This work |
| WT_P<sub>CMV</sub>_CggR_pCggRO | Genomic; Plasmid | P<sub>CMV</sub> and CggR | pCggRO | Wildtype strain containing the CggR under the control of P<sub>CMV</sub> integrated in the genome and the pCggRO reporter plasmid for biosensor analysis | This work |
| WT_P<sub>CMV</sub>_CggR_pYCplac33 | Genomic; Plasmid | P<sub>CMV</sub> and CggR | pYCplac33 | Wildtype strain containing the CggR under the control of P<sub>CMV</sub> integrated in the genome and containing the YCplac33 plasmid for fluorescence background correction | This work |
| WT_P<sub>TEFmut2</sub>_CggR_pCggRO | Genomic; Plasmid | P<sub>TEFmut2</sub> and CggR | pCggRO | Wildtype strain containing the CggR under the control of P<sub>TEFmut2</sub> integrated in the genome and the pCggRO reporter plasmid for biosensor analysis | This work |
| WT_P<sub>TEFmut2</sub>_CggR_pYCplac33 | Genomic; Plasmid | P<sub>TEFmut2</sub> and CggR | pYCplac33 | Wildtype strain containing the CggR under the control of P<sub>TEFmut2</sub> integrated in the genome and containing the YCplac33 plasmid for fluorescence background correction | This work |
| WT_P<sub>TEFmut7</sub>_CggR_pCggRO | Genomic; Plasmid | P<sub>TEFmut7</sub> and CggR | pCggRO | Wildtype strain containing the CggR under the control of P<sub>TEFmut7</sub> integrated in the genome and the pCggRO reporter plasmid for biosensor analysis | This work |
| WT_P<sub>TEFmut7</sub>_CggR_pYCplac33 | Genomic; Plasmid | P<sub>TEFmut7</sub> and CggR | pYCplac33 | Wildtype strain containing the CggR under the control of P<sub>TEFmut7</sub> integrated in the genome and containing the YCplac33 plasmid for fluorescence background correction | This work |
| WT_P<sub>TEFmut2</sub>_R250ACggR_pCggRO | Genomic; Plasmid | P<sub>TEFmut2</sub> and R250ACggR | pCggRO | Wildtype strain containing the mutant R250ACggR under the control of P<sub>TEFmut2</sub> integrated in the genome and the pCggRO reporter plasmid for biosensor analysis | This work |
| WT_P<sub>TEFmut2</sub>_R250ACggR_pYCplac33 | Genomic; Plasmid | P<sub>TEFmut2</sub> and R250ACggR | pYCplac33 | Wildtype strain containing the mutant R250ACggR under the control of P<sub>TEFmut2</sub> integrated in the genome and containing the YCplac33 plasmid for fluorescence background correction | This work |
| WT_P<sub>TEFmut7</sub>_R250ACggR_pCggRO | Genomic; Plasmid | P<sub>TEFmut7</sub> and R250ACggR insertion | pCggRO | Wildtype strain containing the mutant R250ACggR under the control of P<sub>TEFmut7</sub> integrated in the genome and the pCggRO reporter plasmid for biosensor analysis | This work |
| WT_P<sub>TEFmut7</sub>_R250ACggR_pYCplac33 | Genomic; Plasmid | P<sub>TEFmut7</sub> and R250ACggR | pYCplac33 | Wildtype strain containing the mutant R250ACggR under the control of P<sub>TEFmut7</sub> integrated in the genome and containing the YCplac33 | This work |
|  |  |  |  | plasmid for fluorescence background correction |  |
| TM6_pCggRO | Plasmid | *N.A.* | pCggRO | TM6 strain containing the pCggRO reporter plasmid for control of the cis-regulatory dependent YFP/mCherry expression | This work |
| TM6_YCplac33 | Plasmid | *N.A.* | pYCplac33 | TM6 strain containing the YCplac33 plasmid for fluorescence background correction | This work |
| TM6_P<sub>CMV</sub>_CggR_pCggRO | Genomic;<br>Plasmid | P<sub>CMV</sub> and<br>CggR | pCggRO | TM6 strain containing the CggR under the control of P<sub>CMV</sub> integrated in the genome and the pCggRO reporter plasmid for biosensor analysis | This work |
| TM6_P<sub>CMV</sub>_CggR_pYCplac33 | Genomic;<br>Plasmid | P<sub>CMV</sub> and<br>CggR | pYCplac33 | TM6 strain containing the CggR under the control of P<sub>CMV</sub> integrated in the genome and containing the YCplac33 plasmid for fluorescence background correction | This work |
| TM6_P<sub>TEFmut2</sub>_CggR_pCggRO | Genomic;<br>Plasmid | P<sub>TEFmut2</sub> and<br>CggR | pCggRO | TM6 strain containing the CggR under the control of P<sub>TEFmut2</sub> integrated in the genome and the pCggRO reporter plasmid for biosensor analysis | This work |
| TM6_P<sub>TEFmut2</sub>_CggR_pYCplac33 | Genomic;<br>Plasmid | P<sub>TEFmut2</sub> and<br>CggR | pYCplac33 | TM6 strain containing the CggR under the control of P<sub>TEFmut2</sub> integrated in the genome and containing the YCplac33 plasmid for fluorescence background correction | This work |
| TM6_P<sub>TEFmut7</sub>_CggR_pCggRO | Genomic;<br>Plasmid | P<sub>TEFmut7</sub> and<br>CggR | pCggRO | Wildtype strain containing the CggR under the control of P<sub>TEFmut7</sub> integrated in the genome and the pCggRO reporter plasmid for biosensor analysis | This work |
| TM6_P<sub>TEFmut7</sub>_CggR_pYCplac33 | Genomic;<br>Plasmid | P<sub>TEFmut7</sub> and<br>CggR | pYCplac33 | TM6 strain containing the CggR under the control of P<sub>TEFmut7</sub> integrated in the genome and containing the YCplac33 plasmid for fluorescence background correction | This work |
| TM6_P<sub>TEFmut2</sub>_R250ACggR_pCggRO | Genomic;<br>Plasmid | P<sub>TEFmut2</sub> and<br>R250ACggR | pCggRO | TM6 strain containing the mutant R250ACggR under the control of P<sub>TEFmut2</sub> integrated in the genome and the pCggRO reporter plasmid for biosensor analysis | This work |
| TM6_P<sub>TEFmut2</sub>_R250ACggR_pYCplac33 | Genomic;<br>Plasmid | P<sub>TEFmut2</sub> and<br>R250ACggR | pYCplac33 | TM6 strain containing the mutant R250ACggR under the control of P<sub>TEFmut2</sub> integrated in the genome and containing the YCplac33 plasmid for fluorescence background correction | This work |
| TM6_P<sub>TEFmut7</sub>_R250ACggR_pCggRO | Genomic;<br>Plasmid | P<sub>TEFmut7</sub> and<br>R250ACggR | pCggRO | TM6 strain containing the mutant R250ACggR under the control of P<sub>TEFmut7</sub> integrated in the genome and the pCggRO reporter plasmid for biosensor analysis | This work |
| TM6_P<sub>TEFmut7</sub>_R250ACggR_pYCplac33 | Genomic;<br>Plasmid | P<sub>TEFmut7</sub> and<br>R250ACggR | pYCplac33 | TM6 strain containing the mutant R250ACggR under the control of P<sub>TEFmut7</sub> integrated in the genome and containing the YCplac33 plasmid for fluorescence background correction | This work |

**Supplementary Table 7.**
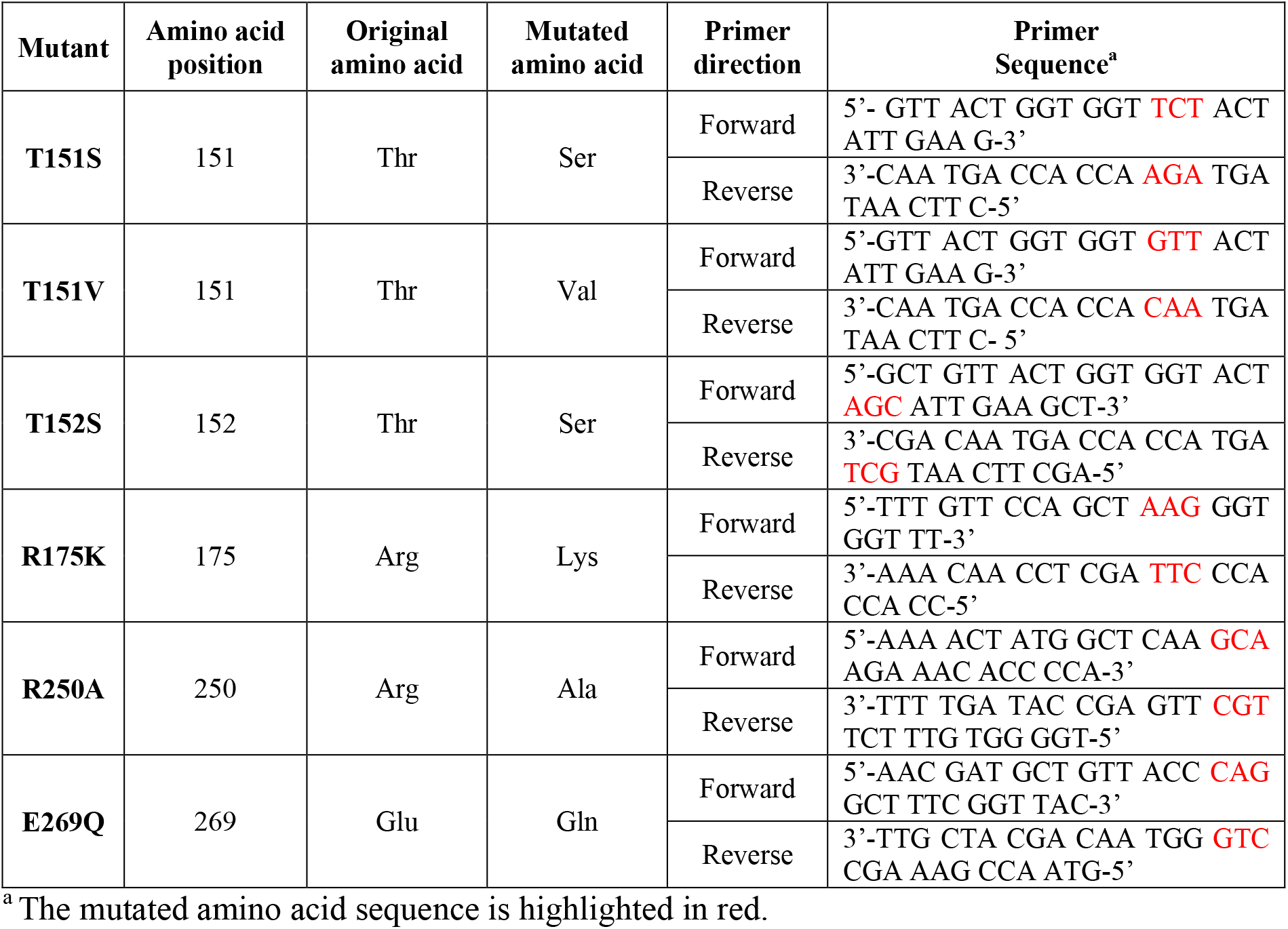
Primers used for CggR site-directed mutagenesis.

**Supplementary Table 8.**
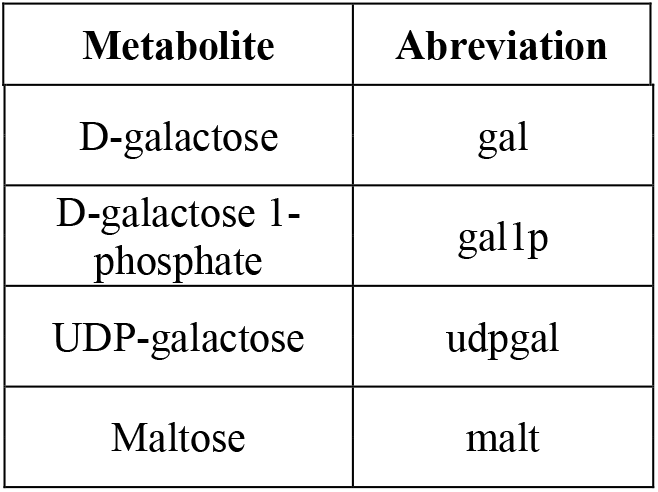
Metabolites that were added to an existing thermodynamic-metabolic network model^43^, which we used here to estimate the intracellular metabolic fluxes.

| Metabolite | Abreviation |
| --- | --- |
| D-galactose | gal |
| D-galactose 1-phosphate | gal1p |
| UDP-galactose | udpgal |
| Maltose | malt |

**Supplementary Table 9.**
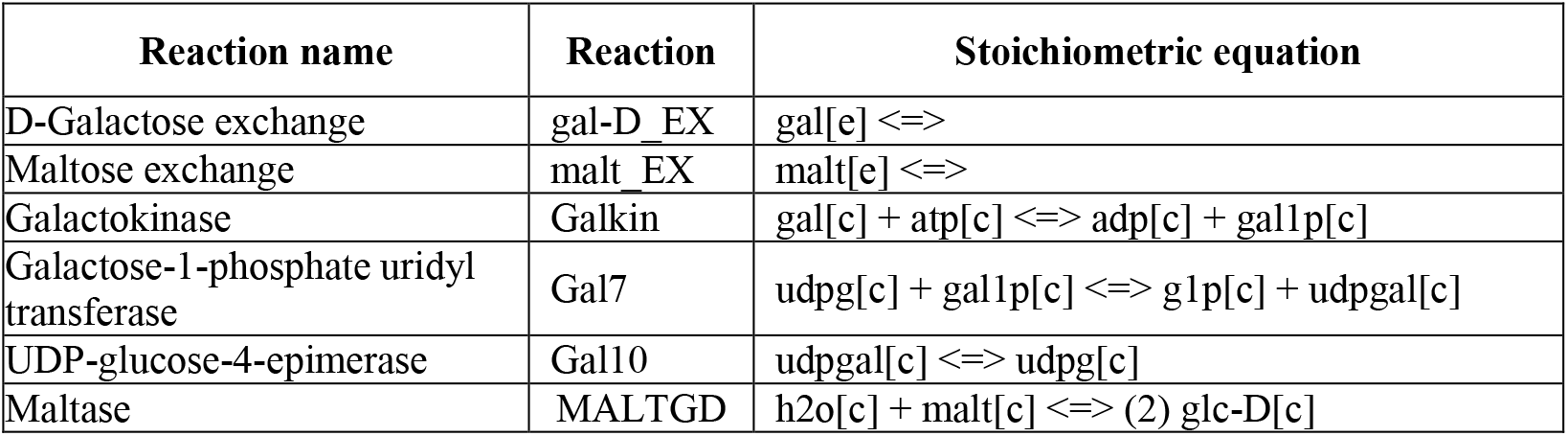
Additional reaction of the metabolic network model. Reactions that were added to an existing thermodynamic-metabolic network model^43^, which we used here to estimate the intracellular metabolic fluxes.

**Supplementary Table 10.**
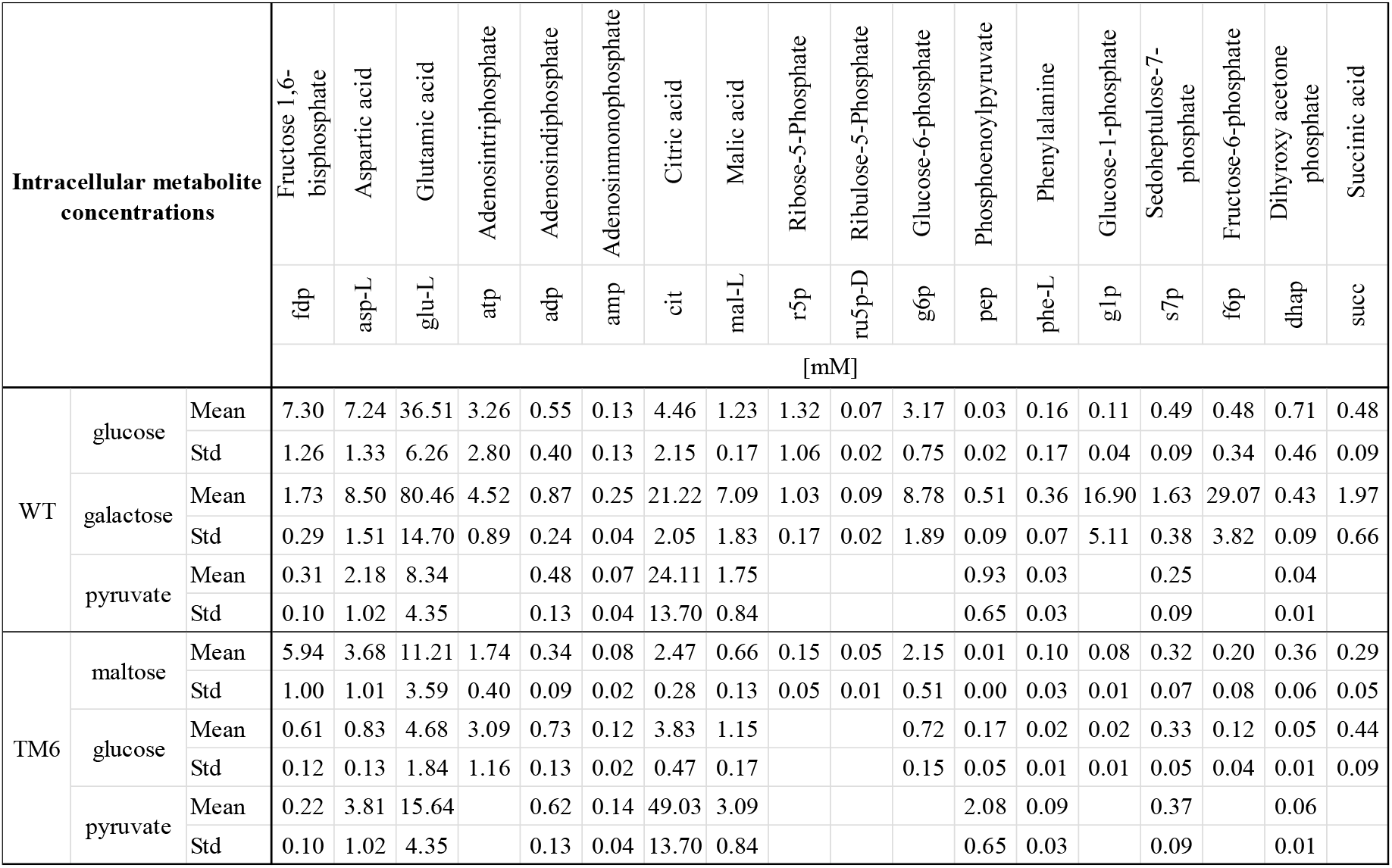
Intracellular metabolite concentrations of S. cerevisiae WT and TM6. The mean (in mM) and standard deviation were calculated from three biological replicates, where each of the biological replicates were sampled three times. For the pyruvate condition, the mean values for WT and TM6 were determined from the values obtained for each strain, but the standard deviations were calculated from the combined samples of WT and TM6.

**Supplementary Table 11.** Physiological parameters used for the intracellular metabolic flux prediction.

|  |  | Parameter | Unit | Mean | Standard deviation |
| --- | --- | --- | --- | --- | --- |
| Wildtype | glucose | $\mu$ [h<sup>-1</sup>] | [h<sup>-1</sup>] | 0.3246 | 0.0038 |
|  |  | Y<sub>x.s</sub> | [g g<sup>-1</sup>] | 0.1149 | 0.0029 |
|  |  | Y<sub>Pyr.s</sub> | [g g<sup>-1</sup>] | 0.0055 | 0.0000 |
|  |  | Y<sub>Gly.s</sub> | [g g<sup>-1</sup>] | 0.0388 | 0.0016 |
|  |  | Y<sub>Ac.s</sub> | [g g<sup>-1</sup>] | 0.0093 | 0.0018 |
|  |  | Y<sub>Etoh.s</sub> | [g g<sup>-1</sup>] | 0.3730 | 0.0034 |
| | galactose | $\mu$ | [h<sup>-1</sup>] | 0.3228 | 0.0130 |
|  |  | Y<sub>x.s</sub> | [g g<sup>-1</sup>] | 0.3759 | 0.0180 |
|  |  | Y<sub>Pyr.s</sub> | [g g<sup>-1</sup>] | 0.0057 | 0.0005 |
|  |  | Y<sub>Gly.s</sub> | [g g<sup>-1</sup>] | 0.0141 | 0.0046 |
|  |  | Y<sub>Ac.s</sub> | [g g<sup>-1</sup>] | 0.0319 | 0.0054 |
|  |  | Y<sub>Etoh.s</sub> | [g g<sup>-1</sup>] | 0.1425 | 0.0160 |
| | pyruvate | $\mu$ | [h<sup>-1</sup>] | 0.0630 | 0.0042 |
|  |  | Y<sub>x.s</sub> | [g g<sup>-1</sup>] | 0.1372 | 0.0086 |
|  |  | Y<sub>Pyr.s</sub> | [g g<sup>-1</sup>] |  |  |
|  |  | Y<sub>Gly.s</sub> | [g g<sup>-1</sup>] |  |  |
|  |  | Y<sub>Ac.s</sub> | [g g<sup>-1</sup>] |  |  |
|  |  | Y<sub>Etoh.s</sub> | [g g<sup>-1</sup>] |  |  |
| TM6 | maltose | $\mu$ | [h<sup>-1</sup>] | 0.3005 | 0.0047 |
|  |  | Y<sub>x.s</sub> | [g g<sup>-1</sup>] | 0.1425 | 0.0022 |
|  |  | Y<sub>Pyr.s</sub> | [g g<sup>-1</sup>] | 0.0033 | 0.0001 |
|  |  | Y<sub>Gly.s</sub> | [g g<sup>-1</sup>] | 0.0089 | 0.0012 |
|  |  | Y<sub>Ac.s</sub> | [g g<sup>-1</sup>] | 0.0164 | 0.0017 |
|  |  | Y<sub>Etoh.s</sub> | [g g<sup>-1</sup>] | 0.3388 | 0.0081 |
| | glucose | $\mu$ | [h<sup>-1</sup>] | 0.2715 | 0.0053 |
|  |  | Y<sub>x.s</sub> | [g g<sup>-1</sup>] | 0.7063 | 0.0280 |
|  |  | Y<sub>Pyr.s</sub> | [g g<sup>-1</sup>] | 0.0006 | 0.0003 |
|  |  | Y<sub>Gly.s</sub> | [g g<sup>-1</sup>] | 0.0015 | 0.0110 |
|  |  | Y<sub>Ac.s</sub> | [g g<sup>-1</sup>] | 0.0060 | 0.0130 |
|  |  | Y<sub>Etoh.s</sub> | [g g<sup>-1</sup>] | 0.0472 | 0.0200 |
| | pyruvate | $\mu$ | [h<sup>-1</sup>] | 0.0765 | 0.0070 |
|  |  | Y<sub>x.s</sub> | [g g<sup>-1</sup>] | 0.3804 | 0.0220 |
|  |  | Y<sub>Pyr.s</sub> | [g g<sup>-1</sup>] |  |  |
|  |  | Y<sub>Gly.s</sub> | [g g<sup>-1</sup>] |  |  |
|  |  | Y<sub>Ac.s</sub> | [g g<sup>-1</sup>] |  |  |
|  |  | Y<sub>Etoh.s</sub> | [g g<sup>-1</sup>] |  |  |

**Supplementary Figure 1.**
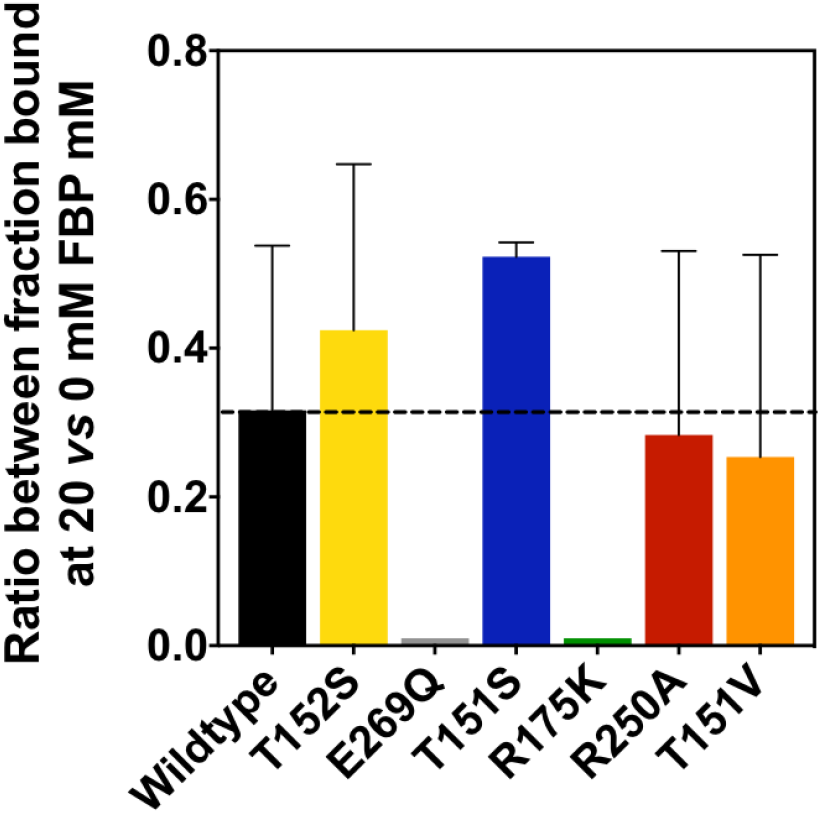
The ratio of CggR bound to DNA at 20 mM FBP *versus* the one at 0 mM FBP represents the FBP-dependent modulation of the CggR-DNA-binding. Error bars correspond to the error calculated from the ratio of the standard deviation of at least three replicates of 0 and 20mM FBP.

**Supplementary Figure 2.**
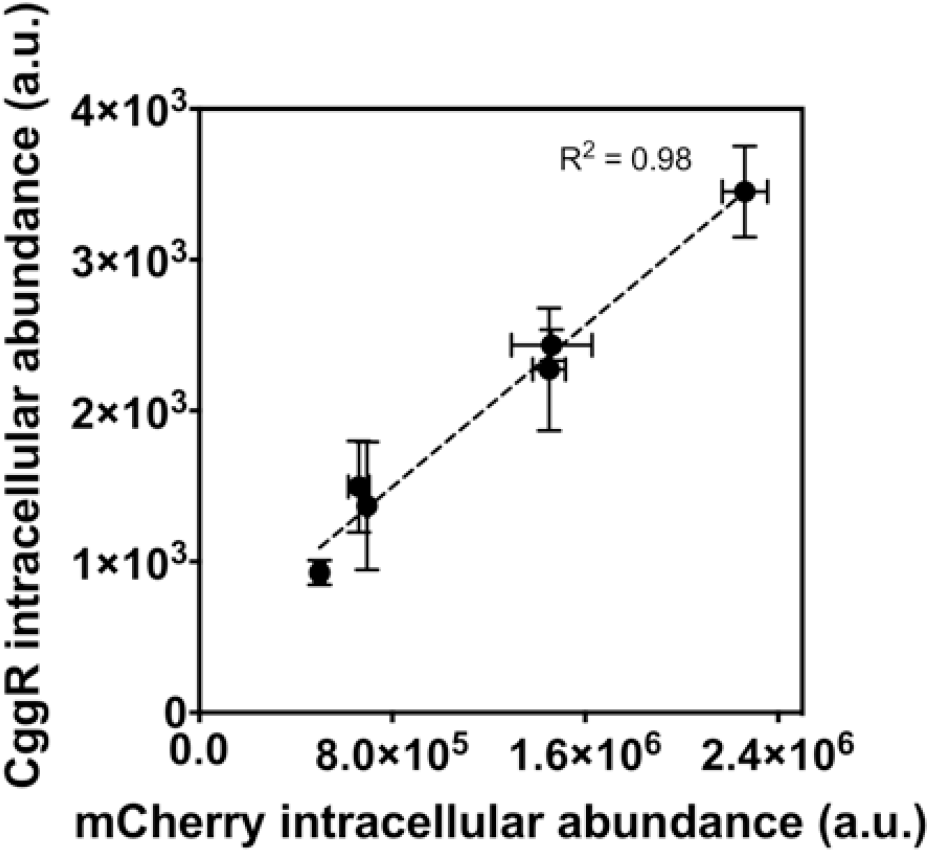
mCherry intracellular levels (abundance) linearly correlate with CggR intracellular levels in wildtype (WT) and TM6 strains. mCherry expression was driven by the P_TEFmut8_ and CggR expression was driven by the P_TEFmut7_ promoter. Error bars indicate the standard deviation of three independent replicates.

**Supplementary Figure 3.**
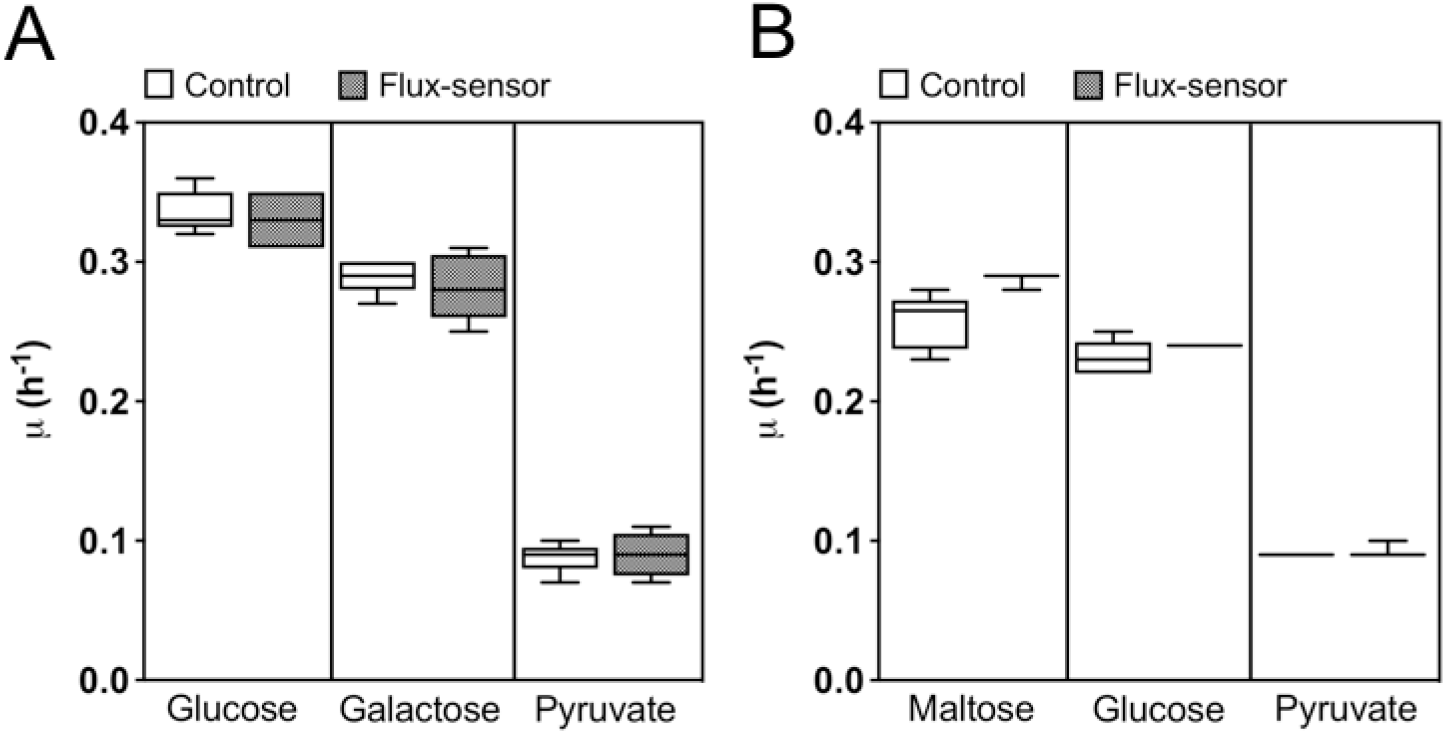
Cellular growth rate is not affected by the expression of the flux-sensor construct in WT (A) and TM6 (B) cells. Error bars represent the standard deviation of at least three replicate experiments.

**Supplementary Figure 4.**
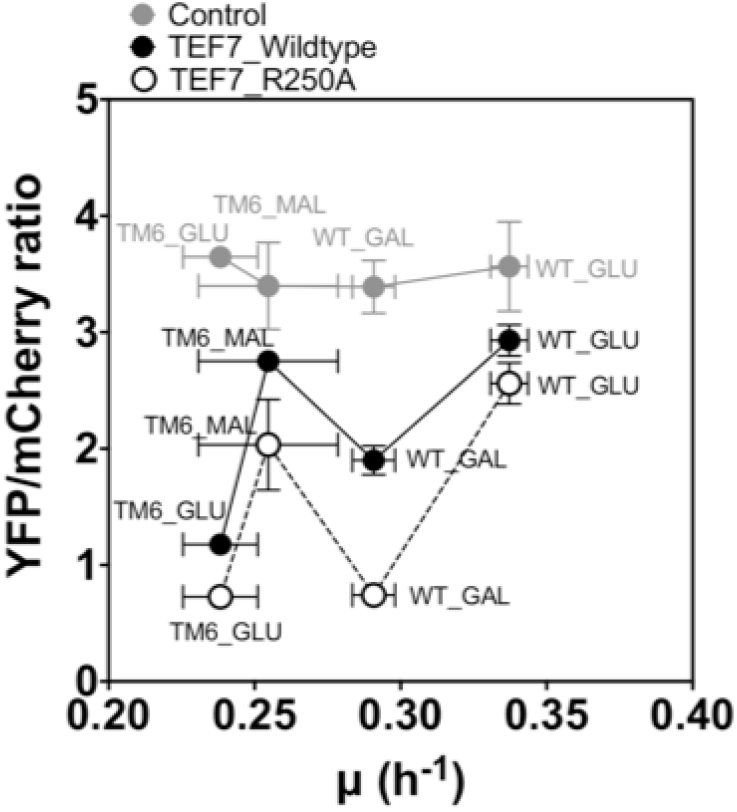
The flux-sensor output shows no correlation with the cellular growth rate. Error bars represent the standard deviation of at least three replicate experiments.

**Supplementary Figure 5.**
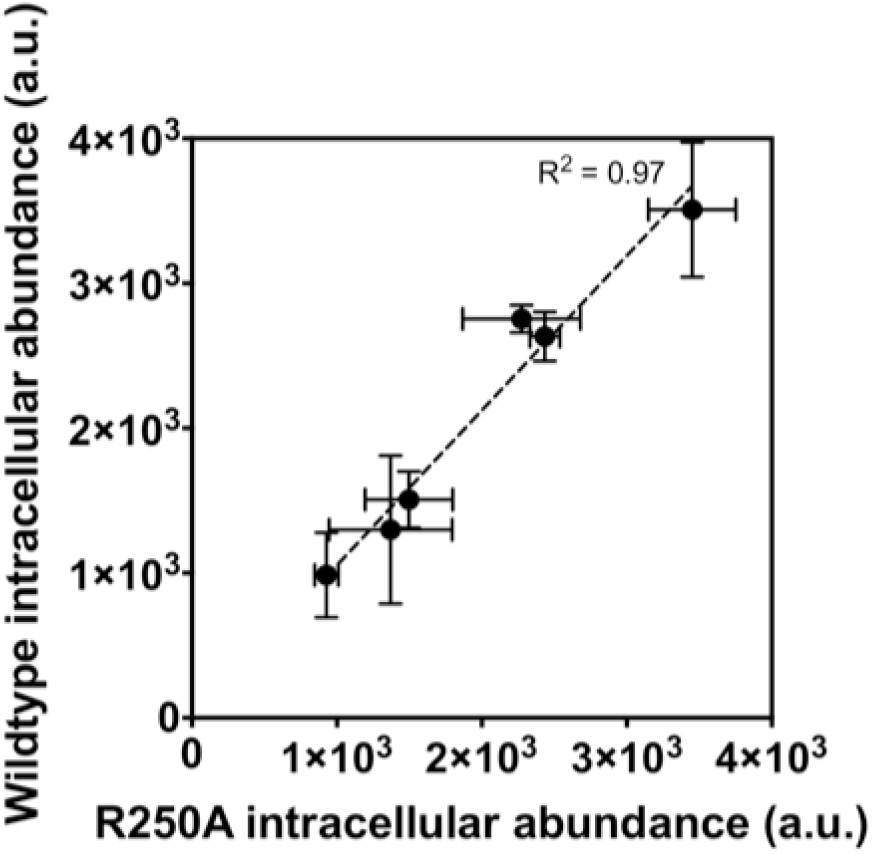
R250A intracellular levels (abundance) linearly correlate with wildtype CggR intracellular levels in WT and TM6 strains. Both CggR variants were expressed by the P_TEFmut7_. Error bars indicate the standard deviation of three independent replicates.

**Supplementary Figure 6.**
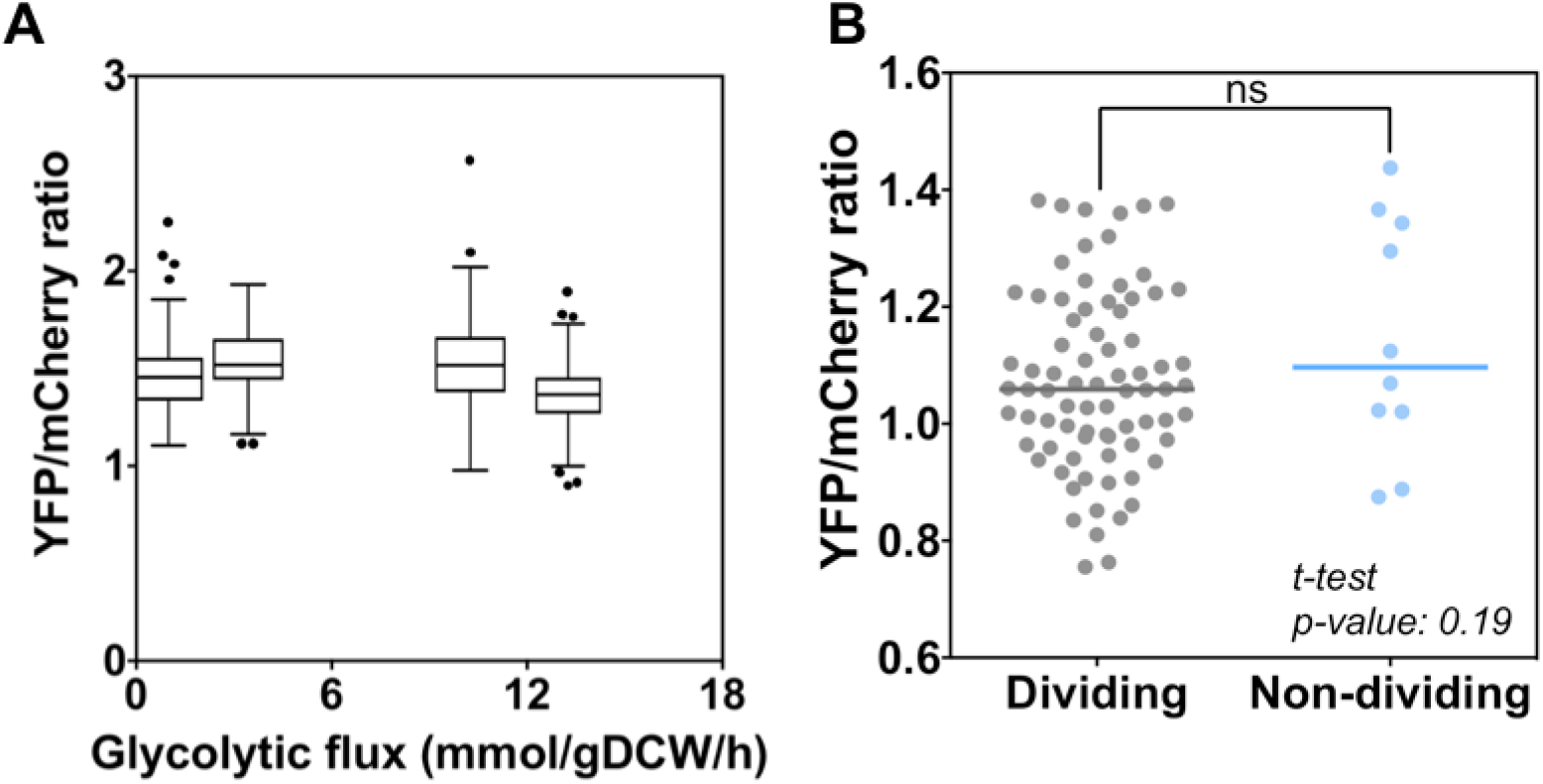
Unregulated controls of the glycolytic flux sensor. (**A**) Tukey boxplots showing the YFP/mCherry ratio of individual cells measured by microscopy for unregulated control strains that don’t express the CggR repressor, cultured in different conditions. At least 35 cells were analyzed in each condition. (**B**) YFP/mCherry ratio measured by microscopy in high-flux (dividing) versus low-flux (non-dividing) TM6 unregulated control cells on 10 gL^-1^ glucose minimal medium.

**Supplementary Figure 7.**
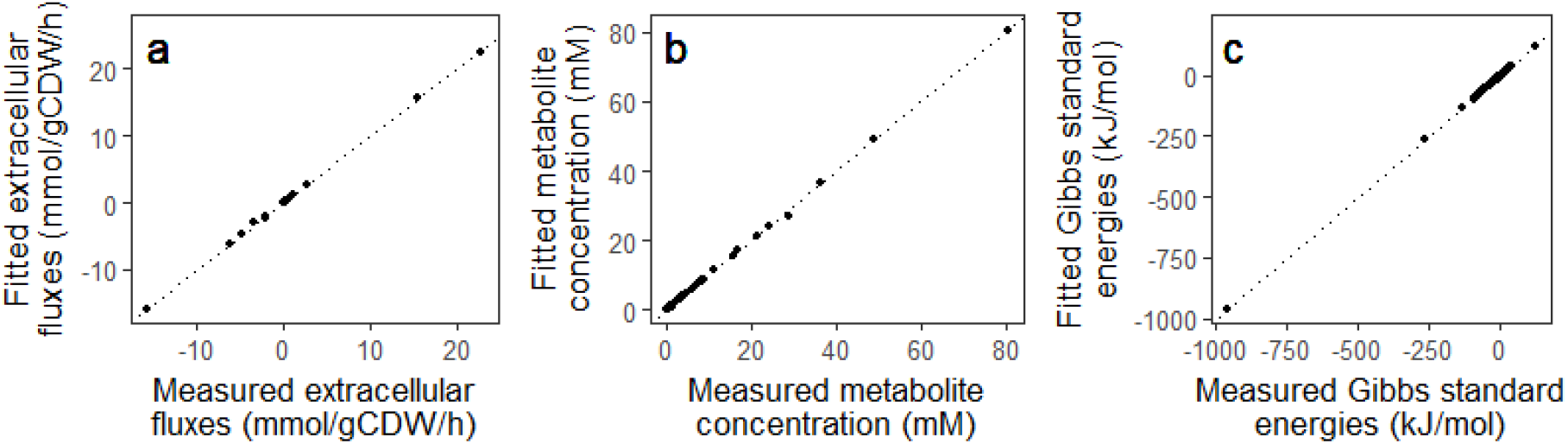
Results of the regression analysis for the six growth conditions of wildtype and TM6 strain. Fitted values from the regression analysis versus measured values; (a) extracellular rates; (b) intracellular metabolite concentrations, and (c) standard Gibbs energies of reactions.

